# Benzo(a)pyrene degradation induces coordinated antioxidant and detoxification responses in the marine yeast *Debaryomyces hansenii*

**DOI:** 10.1101/2025.07.31.667905

**Authors:** Francisco Padilla-Garfias, Minerva Georgina AraizaVillanueva, Martha Calahorra, Norma Silvia Sánchez, Antonio Peña

## Abstract

Polycyclic aromatic hydrocarbons (PAHs), such as benzo(a)pyrene (BaP), are persistent environmental pollutants recognized for their high toxicity and resistance to microbial degradation. In the search for efficient and promising organisms in mycoremediation, the marine yeast *Debaryomyces hansenii* has emerged as a compelling candidate, due to its ability to thrive under a variety of stressful conditions. This study presents the first integrated molecular and biochemical characterization of BaP degradation in a marine yeast, providing mechanistic insights into the detoxification and antioxidant responses involved. In this study, we explored *D. hansenii’*s capacity to degrade BaP and characterized the associated molecular and biochemical responses induced by this compound. Our findings demonstrate that *D. hansenii* can tolerate BaP concentrations up to 100 ppm without compromising cell viability and is capable of degrading nearly 70% of the compound within six days. The degradation process appears to be enzymatically mediated, primarily involving cytochrome P450 (CYP), epoxide hydrolase (EH), and glutathione S-transferase (GST), enzymes typically associated with xenobiotic metabolism, and reactive oxygen species (ROS) generation. BaP exposure resulted in pronounced oxidative stress, evidenced by elevated ROS levels, lipid peroxidation, and protein carbonylation. However, *D. hansenii* activated a robust antioxidant defense, including superoxide dismutase (SOD), catalase (CAT), glutathione peroxidase (GPx), and regulation of the glutathione redox system. Altogether, our data unveil a tightly coordinated cellular strategy that combines oxidation, conjugation, and detoxification pathways to counteract BaP-induced toxicity and sustain redox homeostasis. These insights position *D. hansenii* as a promising and metabolically adaptive organism for bioremediation of PAH-contaminated environments.

## 1. Introduction

Benzo(a)pyrene (BaP) is a five-ring polycyclic aromatic hydrocarbon (PAH) that originates from both natural events (such as volcanic activity) and human-driven processes (including fossil fuel combustion and tobacco use). It has been widely detected in industrial waste products such as coal residues, petroleum sludge, asphalt, and cigarette smoke (Verma et al. 2012). BaP has been found to possess a high degree of chemical stability, a property that contributes to its notable environmental persistence. This characteristic renders BaP resistant to natural degradation processes and predisposes it to accumulation in soils and aquatic sediments (Hattemer-Frey and Travis 1991; Chung et al. 2011). As a result, BaP remains one of the most persistent and ubiquitous toxicants in both terrestrial and aquatic ecosystems (Abdel-Shafy and Mansour 2016).

Human exposure to BaP has risen in recent decades due to the increasing presence of emissions from combustion processes and the consumption of contaminated food products (Bukowska et al. 2022). Once introduced into the body, BaP can be absorbed through multiple pathways, including inhalation, ingestion, and dermal contact (Verma et al. 2012; Bukowska et al. 2022). After internalization, BaP is metabolized by cytochrome P450 (CYP) enzymes, resulting in the formation of highly reactive intermediates (the most important is B(a)P-7,8-diol-9,10-epoxide, also called BPDE) that exhibit carcinogenic, mutagenic, and teratogenic properties (Verma et al. 2012; Ostrem Loss and Yu 2018; Ostrem Loss et al. 2019; Bukowska et al. 2022). Beyond these well-known effects, BaP has also been implicated in neurotoxicity, epigenetic alterations, and reproductive toxicity, underscoring its broad impact on both human health and ecological integrity (Verma et al. 2012; Ghosal et al. 2016; Bukowska et al. 2022).

Due to BaP’s persistence and toxicological burden, there is an increasing interest in identifying sustainable and biologically driven remediation strategies. Among these, mycoremediation (the use of fungi to degrade or transform organic pollutants) has gained attention for its ecological compatibility and enzymatic versatility (Deshmukh et al. 2016; Park and Choi 2020). Although conventional approaches often rely on biochemical assays to measure fungal degradation capacity, the underlying metabolic and regulatory mechanisms remain poorly understood (Clemente et al. 2001; Cerniglia and Sutherland 2010; Nayak et al. 2013; Aranda 2016; Park and Choi 2020). Research in this field has predominantly focused on filamentous fungi from the Ascomycota and Basidiomycota phyla (Harms et al. 2011; Park and Choi 2020). In contrast, yeasts, offering rapid growth, ease of genetic manipulation, and notable stress tolerance, have received limited attention, despite their considerable biotechnological promise (Aranda 2016; Padilla-Garfias et al. 2024).

In Ascomycota fungi, including yeasts, the degradation of PAHs typically follows a three-phase metabolic pathway. The process begins with the oxidation of aromatic rings, typically catalyzed by CYP, their NADPH-dependent reductases (CPR), and epoxide hydrolases (EH), which initiate the oxidation of complex hydrocarbon structures. Secondly, the reactive intermediates generated are conjugated to functional groups (such as glutathione, sulfate, methyl, or glucose) by transferases, particularly glutathione S-transferases (GST) in the case of glutathione as the utilized functional group. These conjugates are subsequently either actively exported from the cell or stored into the vacuole (Verdin et al. 2005; Fayeulle et al. 2014; Aranda 2016). This integrated enzymatic system, collectively referred to as the xenome, constitutes the cellular machinery responsible for detoxifying xenobiotics (Morel et al. 2013; Aranda 2016; Padilla-Garfias et al. 2021; Padilla-Garfias et al. 2024). A comprehensive understanding of the molecular regulation and functional dynamics of this system is essential for refining fungal biotechnological applications in environmental remediation.

A central challenge in BaP degradation is the oxidative stress induction that is generated during the initial oxidation phase. Reactive oxygen species (ROS), which are formed as metabolic by-products, have been shown to damage cellular macromolecules, including lipids, proteins, and nucleic acids (Jamieson 1998; Ikner and Shiozaki 2005; Herrero et al. 2008; Li et al. 2009; Yaakoub et al. 2022). To mitigate these negative effects, it is imperative that fungal cells activate antioxidant defense mechanisms, including enzymes such as superoxide dismutase (SOD), catalase (CAT), and glutathione peroxidase (GPx), which collectively mitigate oxidative damage. Central to this system is reduced glutathione (GSH), a multifunctional tripeptide that plays a pivotal role in neutralizing free radicals and maintaining intracellular redox balance (Emri et al. 1997; Livingstone 2001; Li et al. 2009; Schmacht et al. 2017; Yaakoub et al. 2022; Padilla-Garfias et al. 2024).

In this context, the marine yeast *Debaryomyces hansenii* has attracted considerable interest due to its unique adaptability to extreme environmental conditions, including high salinity, tolerance to heavy metals, and its metabolic versatility (Nobre et al. 1999; Breuer and Harms 2006; Prista and Loureiro-Dias 2007; Sánchez et al. 2008; Aggarwal and Mondal 2009; Navarrete et al. 2009; Gumá-Cintrón et al. 2015; Prista et al. 2016; Sánchez et al. 2020; Navarrete et al. 2022). In previous work, our group demonstrated that *D. hansenii* can degrade BaP via a CYP enzyme encoded by the *DhDIT2* gene (Padilla-Garfias et al. 2022). However, that study was primarily focused on gene identification and did not explore the cellular responses, biochemical pathways, or oxidative stress mechanisms associated with BaP degradation. In the present work, we seek to bridge this gap by providing a comprehensive characterization of the functional response of *D. hansenii* upon BaP exposure. These insights reinforce the potential of this yeast as a robust candidate for mycoremediation, particularly in marine and coastal environments impacted by PAH contamination.

This study proposes that BaP exposure elicits a coordinated response in *D. hansenii*, engaging both detoxification and antioxidant defense mechanisms that collectively support cellular viability and metabolic resilience under this chemical stress. Despite evidence of BaP degradation by *D. hansenii*, the cellular and biochemical responses that enable this process remained unclear; to evaluate this proposal, we implemented an integrative experimental strategy combining physiological, biochemical, and molecular assays to assess growth performance, BaP degradation efficiency, and activation of antioxidant pathways. Our findings provide new understanding into the adaptive landscape of *D. hansenii* in response to PAH exposure and highlight its promise as a robust, stress-tolerant organism for biotechnological applications in the remediation of persistent environmental pollutants like BaP.

## 2. Materials and methods

### 2.1 Strain and cultures

*Debaryomyces hansenii* Y7426 strain (United States Department of Agriculture, Peoria, Illinois, USA) was maintained on solid YNBG medium (0.67% yeast nitrogen base with amino acids, ammonium sulphate, supplemented with 2% glucose, and 2% agar). Cultures were refreshed monthly and routinely grown overnight in 250 mL of YNBG in 500 mL Erlenmeyer flasks at 28 °C with orbital shaking at 250 revolutions per minute (rpm) for 24 hours. All experimental cultures were initiated at an optical density at 600 nm (OD_600_= 0.1).

### 2.2 Growth and survival assays

All growth and survival assays, unless otherwise indicated, were performed under aseptic conditions using three independent biological replicates, each measured in technical duplicates. To assess BaP tolerance, *D. hansenii* cells were pre-cultured for 24 hours in YNBG, washed twice with sterile water, and resuspended at an OD_600_= 1.0. 10-fold serial dilutions were prepared in sterile water in 96-well plates, and 5 µL of each dilution were spotted onto the surface of YNBG (control) and YNB + 100 ppm BaP (experimental), using an aluminum multi-pin replicator. Plates were incubated at 28 °C and monitored daily for six days.

For growth curves analysis, a 24-hour pre-culture was adjusted to an OD_600_ of 0.03, using a Beckman DU 650 spectrophotometer and inoculated into the respective media (YNBG or YNB + 100 ppm BaP). Cultures were incubated at 28 °C, and optical density was recorded hourly for six days using a Bioscreen C automated plate reader (with an initial OD_600_ baseline of approximately 0.2).

Dry weight determinations were performed over 6 days; pre-cultures grown in YNBG were used to inoculate 50 mL of fresh YNBG or YNB + 100 ppm BaP, adjusting to OD_600_ = 0.1. At 24-hours intervals, 1.0 mL aliquots were collected and filtered through pre-weighed 0.22 µm filter papers using a vacuum filtration system (Sampling Manifold 1225, Merck Millipore) to remove the culture medium. The filters were then washed twice with sterile water, and excess liquid was removed under vacuum. Filter papers were subsequently transferred to aluminum plates and dried at 95 °C until reaching a constant weight. Final weights were recorded using an analytical balance.

### 2.3 BaP degradation assay

BaP degradation was quantified by spectrofluorometry, following the method described by Padilla-Garfias et al. (2022). All assays were conducted in triplicate using three independent biological replicates. Cultures of *D. hansenii* grown for 24 hours in YNBG were used to inoculate 250 mL of YNB + 100 ppm BaP, adjusted to an initial OD_600_ = 0.1. Samples (3.0 mL) were collected at 0, 1, 2, 3, and 6 days and stored at −20 °C for BaP extraction.

Extraction was performed with chloroform, followed by evaporation and resuspension in acetone. Fluorescence was measured using an AMINCO SLM spectrofluorometer (excitation wavelength at 356 nm, emission wavelength at 405 nm), and concentrations were calculated from a standard curve. To distinguish true biodegradation from passive adsorption or abiotic loss, two negative controls were included: a cell-free medium control and a heat-inactivated culture control (95 °C, 15 minutes).

### 2.4 RNA extraction

*D. hansenii* cells were initially grown in 250 mL of YNBG medium at 28 °C with shaking at 250 rpm for 24 hours. Following this pre-culture, 50 mL flasks containing YNBG or YNB + 100 ppm BaP at an OD_600_ = 0.1 were inoculated and incubated under the same conditions for 0, 1, 2, 3, and 6 days.

For RNA extraction, 15 mL samples were collected on day 0 (after pre-culture) and the remaining days (1, 2, 3, and 6). Cells were pelleted by centrifugation for 5 min at 3000 rpm (1750 x g) and resuspended in 1.0 mL of AE buffer (50 mM sodium acetate, 10 mM ethylenediaminetetraacetic acid (EDTA), pH 5.2). Total RNA was extracted using the method described by Schmitt et al. (1990). RNA quality and integrity were assessed by 1% denaturing agarose gel electrophoresis, confirming the presence of intact 28S and 18S rRNA bands.

### 2.5 Gene expression studies: cDNA synthesis and RT-qPCR

Total RNA was treated with DNase I (RQ1 RNase-Free DNase kit, Promega) to eliminate genomic DNA contamination, and reverse transcribed into cDNA using the ImProm-II™ Reverse Transcription System (Promega). Gene expression was subsequently analyzed by RT-qPCR using three independent biological replicates and two technical replicates per sample.

RT-qPCR was performed using specific primers (see Supplementary Table 1) targeting 24 open reading frames (ORFs) found to be up-regulated in NCBI-deposited RNA-Seq (NCBI Gene Expression Omnibus database under accession No. GSE299919) from our previous work (Padilla-Garfias et al. 2025), associated with detoxifying metabolism and genes coding for antioxidant enzymes, searching gene ontologies in MycoCosm (https://mycocosm.jgi.doe.gov/mycocosm/home (Grigoriev et al. 2014)) and added in Supplementary Table 2. Sequences were retrieved from NCBI, and primers were designed using Primer-BLAST, then evaluated for dimer formation and specificity with DINAMelt (http://www.unafold.org/hybrid2.php) (Markham and Zuker 2005). Primers were synthesized at the Molecular Biology Unit of the Institute of Cellular Physiology, UNAM.

qPCR reactions were run on a Rotor-Gene Q thermal cycler (Qiagen) using SYBR Green chemistry (qPCR SyberMaster highROX, Jena Bioscience). Relative gene expression was quantified using the 2⁻^ΔΔCT^ method (Livak and Schmittgen 2001), with YNBG-grown cells as the calibrator. Expression values were normalized to *DhACT1* (ORF DEHA2D05412g), following Sánchez et al. (2020).

### 2.6 Preparation of protein extracts

Cells were cultured in 250 mL YNBG medium and YNB + 100 ppm BaP. Samples were collected on days 0, 1, 2, 3, and 6 and centrifuged 5 min at 3000 rpm (1400 x g) in a Beckman Coulter Avanti J-26 XPI centrifuge. Pellets were resuspended at 1 g/mL (wet weight/volume) in 10 mM 3-(N-morpholino) propanesulfonic acid (MOPS) buffer, pH 7, containing protease inhibitors (cOmplete, Roche). Glass beads (0.45 mm diameter) were added to half-fill the sample volume, and cells were disrupted using a Bead Beater chamber placed on ice, performing six 30-second homogenizing cycles with 2-minute intervals between each. All steps were carried out at 4 °C. Cell debris was removed by centrifugation at 3000 rpm (1750 x g) for 5 minutes in an IEC clinical centrifuge.

### 2.7 Enzymatic activities

Prior to the enzymatic assays, the supernatants from the cell extracts obtained as described in section 2.6 of Materials and Methods were centrifuged at 10,000 rpm (6700 x g) for 10 minutes in an Eppendorf centrifuge 5415C. Protein concentration was then determined using the Lowry method as modified by Markwell in a Beckman DU 650 spectrophotometer (Markwell et al. 1978). All enzymatic measurements were performed using an Aminco DW-2a UV/VIS spectrophotometer (OLIS, On-Line Instrument Systems, Inc. converted to DW2).

The activities of cytochrome P450 (CYP) and NADPH-cytochrome c reductase (CPR) were indirectly estimated by quantifying CPR activity using a commercial assay kit (Cytochrome c Reductase (NADPH) Assay Kit CY0100, Sigma, Saint Louis, MO, USA). Enzyme activity values were normalized to protein content (mg) as described previously (Padilla-Garfias et al. 2022).

Epoxide hydrolase (EH) activity was determined according to Mateo et al. (2023). Briefly, 3 mL of reaction mixture containing 100 mM sodium phosphate buffer (pH 7.0), 2 mM styrene oxide (prepared in dimethylformamide), 2 mM sodium metaperiodate, and 40 µg of protein extract were incubated at 30 °C. Enzymatic activity was monitored by the increase in absorbance at 290 nm over 2 minutes. Specific activity was calculated using the molar extinction coefficient of benzaldehyde (ε = 1.34 mM⁻¹ cm⁻¹).

Glutathione S-transferase (GST) activity was measured following the method of Habig and Jakoby (1981), with slight modifications. The 2 mL reaction mixture contained 100 mM sodium phosphate buffer (pH 6.5), 5 mM reduced glutathione (GSH), 1 mM 1-chloro-2,4-dinitrobenzene (CDNB; prepared in ethanol), and 40 µg of protein extract. The formation of the GSH-CDNB adduct was monitored by the increase in absorbance at 340 nm for 3 minutes at 30 °C. Specific activity was calculated using the molar extinction coefficient of the adduct (ε = 9.6 mM⁻¹ cm⁻¹).

Glutathione reductase (GR) activity was estimated based on the method of Carlberg and Mannervik (1985), with modifications. The assay was performed in a 2 mL mixture containing 100 mM sodium phosphate buffer (pH 7.0), 1 mM oxidized glutathione (GSSG), 0.15 mM NADPH, and 40 μg of protein extract. The decrease in absorbance at 340 nm, due to NADPH oxidation, was recorded for 2 minutes at 30 °C. Specific activity was calculated using the NADPH molar extinction coefficient (ε = 6.2 mM⁻¹ cm⁻¹).

Glutathione peroxidase (GPx) activity was assayed following Gonzalez et al. (2020), with modifications. The 2 mL reaction mixture included 100 mM sodium phosphate buffer (pH 7.0), 1 mM GSH, 0.15 mM NADPH, 1 unit of GR, 1 mM hydrogen peroxide (H_2_O_2_), and 40 μg of protein extract. The decrease in absorbance at 340 nm was monitored for 1 minute at 30 °C. Specific activity was calculated using the NADPH extinction coefficient (ε = 6.2 mM⁻¹ cm⁻¹).

Superoxide dismutase (SOD) activity was measured according to González et al. (2020), with modifications. Reactions were performed in 2 mL mixtures containing 50 mM sodium phosphate buffer (pH 7.0), 0.1 mM EDTA, 13 mM methionine, 75 µM nitroblue tetrazolium (NBT), 2 µM riboflavin, and 40 µg of protein extract. Controls received equivalent volumes of buffer in place of protein. Tubes were exposed to fluorescent light (from a transilluminator UVP) for 10 minutes to initiate the reaction, followed by a 10-minute incubation in the dark. Absorbance was recorded at 560 nm. SOD activity was expressed as the enzyme amount required to inhibit 50% of NBT reduction, using the following formula:

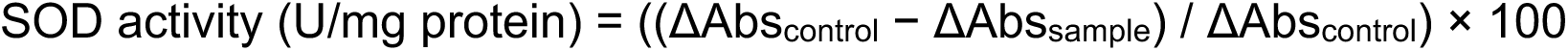

Catalase (CAT) activity was estimated based on Aebi (1984), with modifications. The reaction mixture contained 100 mM sodium phosphate buffer (pH 7.0) and 1 mM H_2_O_2_. The decomposition of H_2_O_2_ was monitored by the decrease in absorbance at 240 nm over 2 minutes at 30 °C. Specific CAT activity was calculated using a calibration curve prepared with H_2_O_2_ concentrations ranging from 1 nM to 10 mM.

### 2.8 Quantification of reduced glutathione (GSH) and oxidized glutathione (GSSG)

For glutathione quantification, the supernatants from cell extracts prepared as described in sections 2.6 and 2.7 of Materials and Methods; concentrations were normalized to the total protein content and expressed as µmol of GSH or GSSG per mg of protein.

Glutathione levels were determined following the method of Griffith (1980). Extracts were deproteinized with 300 mM phosphoric acid, incubated at 4 °C for 20 minutes, and centrifuged at 10,000 rpm (6700 x g) for 10 minutes in an Eppendorf centrifuge 5415C. GSH levels were quantified based on its reaction with DTNB (5,5’-dithiobis-(2-nitrobenzoic acid)) at 412 nm. A standard calibration curve was prepared using serial dilutions of GSH ranging from 0 to 100 µM to calculate sample concentrations.

To measure GSSG, free GSH was first derivatized with 2-vinylpyridine. GSSG was then reduced to GSH using GR and NADPH, and the absorbance of the resulting DTNB-GSH complex was measured at 412 nm. A separate standard curve was generated using GSSG standards (0–10 µM), subjected to the same derivatization and reduction steps.

### 2.9 Quantification of oxidative stress markers

Intracellular accumulation of reactive oxygen species (ROS) was assessed over a six-day period using two fluorescence-based probes with subcellular specificity. Cytoplasmic ROS were detected using 10 µM of H₂DCF-DA (2′,7′-dichlorodihydrofluorescein diacetate), while mitochondrial ROS were evaluated with 5 µM of dihydrorhodamine 123 (DHR123) (Pérez-Gallardo et al. 2013). Cells grown in YNBG or YNB medium supplemented with 100 ppm of BaP were harvested daily and incubated with the respective probes for 30 minutes at 30 °C in the dark. Fluorescence was subsequently measured using a POLARstar Omega microplate reader (BMG Labtech, Germany), with excitation/emission wavelengths of 435/530 nm for H₂DCF-DA and 500/536 nm for DHR123. Fluorescence values were normalized to cell density, determined by OD_600_ measurements on the same device (James et al. 2015).

Lipid peroxidation was monitored daily throughout the six-day period by quantifying malondialdehyde (MDA), a stable end-product of polyunsaturated fatty acid oxidation. Cell extracts were prepared as described in section 2.6 and deproteinized by adding 20% (w/v) trichloroacetic acid (TCA). Samples were incubated on ice for 10 minutes and centrifuged at 10,000 rpm (6700 x g) in an Eppendorf centrifuge 5415C for 10 minutes at 4 °C. The supernatant was then incubated with thiobarbituric acid (TBA) for 60 minutes at 95 °C. After cooling, the absorbance of the MDA-TBA adduct was measured at 532 nm using a Beckman DU 650 UV/VIS spectrophotometer. MDA concentration was calculated using a molar extinction coefficient (ε = 156 mM⁻¹ cm⁻¹) and normalized to the total protein content (Ohkawa et al. 1979).

Oxidative damage to proteins was also assessed daily over the six-day period by quantifying carbonylated protein content in cells grown under the same conditions. Cell extracts were prepared as previously described in sections 2.6 and 2.7 of Materials and Methods. The resulting supernatants were incubated with 5 mM 2,4-dinitrophenylhydrazine (DNPH) for 1 hour at room temperature in the dark, with occasional mixing. Protein hydrazone derivatives were precipitated with 20% TCA, washed thoroughly with an ethanol-acetone solution (1:1, v/v), and resuspended. Absorbance was recorded at 370 nm using a Beckman DU 650 spectrophotometer. Carbonyl content was calculated using a molar extinction coefficient (ε = 22 mM⁻¹ cm⁻¹) and normalized to the total protein content (Levine et al. 1990).

### 2.10 Statistical analysis

Data are presented as mean ± standard deviation from three independent biological replicates, each measured in triplicate. Statistical significance was evaluated by two-way ANOVA or Student’s *t*-test in Prism 10 (GraphPad Software Inc., San Diego, CA, USA), with 95% confidence intervals.

## 3. Results

*D. hansenii* is known for its remarkable resistance to extreme environments, including high salinity, low temperature, heavy metals, alternative carbon sources, linear hydrocarbons, and oxidative stress conditions (Yadav and Loper 1999; Nobre et al. 1999; Breuer and Harms 2006; Prista and Loureiro-Dias 2007; Sánchez et al. 2008; Aggarwal and Mondal 2009; Navarrete et al. 2009; Bonugli-Santos et al. 2015; Gumá-Cintrón et al. 2015; Prista et al. 2016; Ramos-Moreno et al. 2019; Sánchez et al. 2020; Navarrete et al. 2022; de la Fuente-Colmenares et al. 2024), traits that are crucial for its ability to metabolize toxic compounds such as BaP (Cerniglia and Crow 1981; Padilla-Garfias et al. 2022; Padilla-Garfias et al. 2025). Exposure to BaP triggers complex transcriptomic and biochemical responses in this yeast (Gumá-Cintrón et al. 2015; Ostrem Loss et al. 2019; Park and Choi 2020; Martínez-Ávila et al. 2021; Padilla-Garfias et al. 2025). Its capacity to thrive in such adverse settings is supported by an efficient oxidative stress response, including a notable tolerance to H_2_O_2_ and other reactive agents. These features suggest that *D. hansenii* possesses robust detoxification mechanisms to counteract BaP-induced oxidative damage (Navarrete et al. 2009; Segal-Kischinevzky et al. 2011; Michán et al. 2013; Gumá-Cintrón et al. 2015; Ramos-Moreno et al. 2019; de la Fuente-Colmenares et al. 2024).

To investigate and understand the response of *D. hansenii* to BaP exposure, a multi-level experimental approach was employed. This approach integrated growth and degradation assays, gene expression analysis, targeted RT-qPCR, enzyme activity measurements, and quantification of oxidative stress markers.

Tolerance and growth assays revealed that *D. hansenii* was able to grow in the presence of 100 ppm BaP, both in liquid and solid media, also demonstrating the ability to degrade the compound (Figure 1). When cultured with BaP as sole carbon source at 28 °C in solid media, the yeast exhibited growth comparable to the control condition, consistent with previous findings (Padilla-Garfias et al. 2022; Padilla-Garfias et al. 2025) (Figure 1a). Fluorescence-based measurements showed that *D. hansenii* was able to degrade approximately 70% of the BaP within six days (Figure 1b).

**Figure 1.**
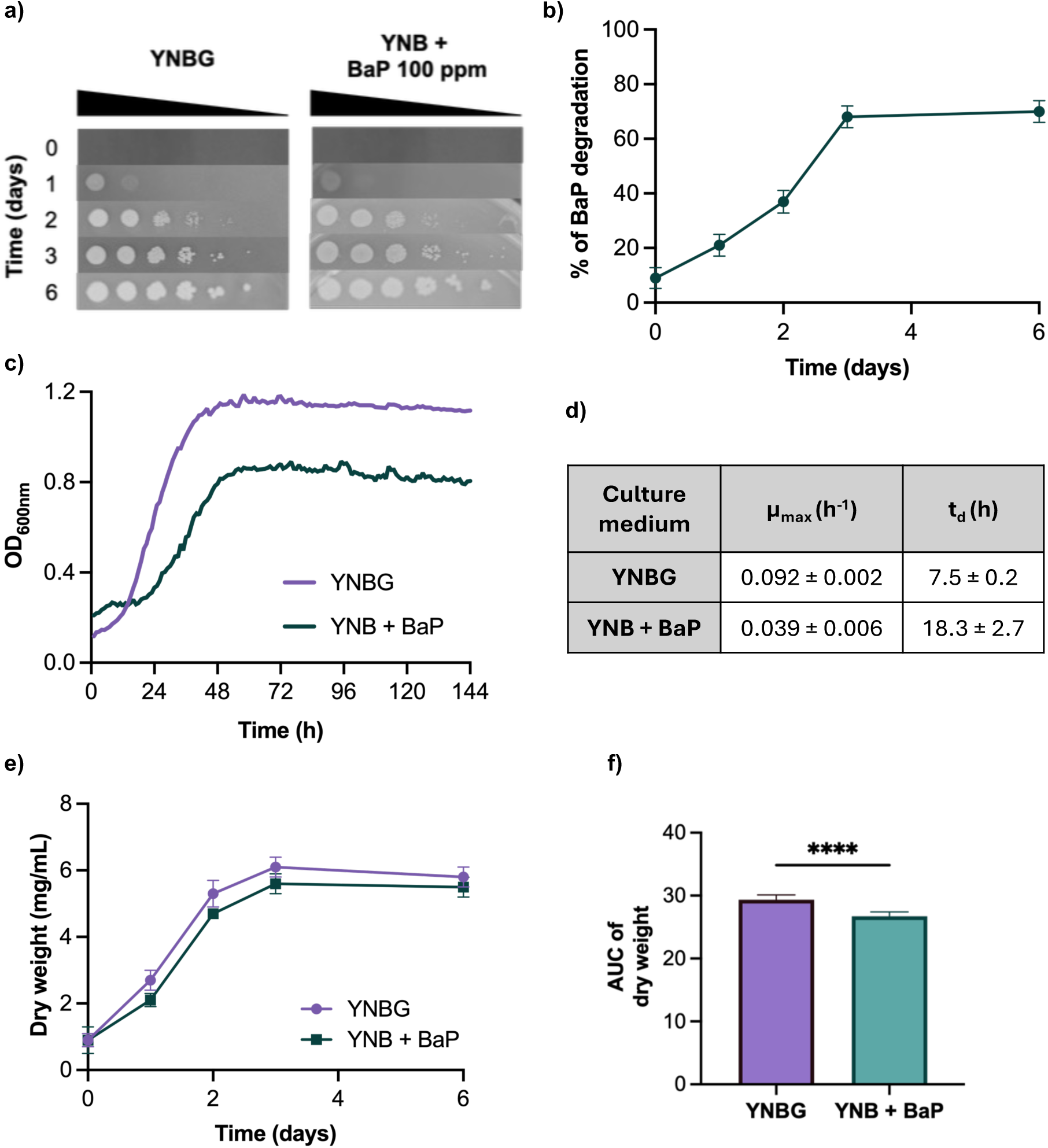
Growth and degradation capacity of *D. hansenii* in response to BaP exposure. a) Spot assay on solid YNBG and YNB + 100 ppm BaP plates assessed by serial dilutions after 6 days of incubation at 28 °C (representative image of n = 9). b) BaP degradation measured by fluorescence-based detection over six days (n = 9, three biological and three technical replicates). c) Growth curves of *D. hansenii* in YNBG and YNB + 100 ppm BaP at 28 °C. Optical density at 600 nm (OD_600_) was recorded every 30 min using a microplate reader (n = 30, three biological and ten technical replicates). d) Specific growth rate (μ_max_) and doubling time (t_d_) calculated from the exponential phase of growth curves shown in c). e) Biomass accumulation over six days, measured as dry weight (n = 18, three biological and six technical replicates). f) Area under the curve (AUC) of biomass production was calculated for statistical comparison. Significance was assessed using two-tailed unpaired Student’s t-test (p < 0.05, ****p < 0.0001).

In liquid culture (Figure 1c), the duplication time increased from 7.5 hours with glucose to 18.3 hours with BaP, reflecting a reduced specific growth rate under these conditions. Although key kinetic parameters were altered (Figure 1d), overall cell density remained lower when *D. hansenii* grew under glucose-deficient conditions. Biomass production, estimated by dry weight over a six-day period, is shown in Figure 1e. A minor decrease in biomass was observed from day 3 onward in glucose-grown cells lacking BaP. Nevertheless, the differences in biomass yield between the conditions were statistically significant after calculating the areas under the curve (AUC) in order to perform the necessary statistical analyses (Figure 1f). Biomass determination (Figure 1e) is critical for quantifying microbial growth and for optimizing biotechnological applications such as mycoremediation.

Initial physiological and metabolic assays were used to confirm cell viability, which are essential for a reliable detoxification process. Once *D. hansenii*’s ability to grow and degrade BaP was established, we analyzed RNA-Seq data obtained on day 3 of growth (NCBI Gene Expression Omnibus database under accession No. GSE299919 (Padilla-Garfias et al. 2025)). This analysis identified 24 ORFs associated with detoxification pathways, glutathione homeostasis, and antioxidant enzymes (Supplementary Table 2). Based on these findings, we designed specific primers (Supplementary Table 1) and performed RT-qPCR on days 0, 1, 2, 3 and 6 of growth under both BaP-exposed and control conditions. The complete RT-qPCR datasets are presented in Supplementary Figure 1, whereas the figures throughout the Results section display heatmap summaries derived from that data.

The degradation of BaP in *D. hansenii* is orchestrated by a group of detoxifying enzymes whose genes are significantly upregulated in response to the contaminant (Padilla-Garfias et al. 2025). Also illustrated in Figures 2a, 2c, and 2e; these ORFs encode enzymes including CYP, CPR, EH, GST. CYP and CPR initiate phase I of xenobiotic metabolism by oxidizing BaP into highly reactive epoxide intermediates (Aranda 2016; Ostrem Loss et al. 2019; Padilla-Garfias et al. 2022; Padilla-Garfias et al. 2024). These epoxides are subsequently converted into less toxic diols by EH during phase II (Marco-Urrea et al. 2015; Aranda 2016). GST then conjugate GSH to the resulting metabolites, facilitating their removal or vacuolar sequestration (Capotorti et al. 2005; Ghosal et al. 2016; Varjani 2017). Enzymatic activity measurements (Figures 2b, 2d, and 2f) revealed a peak in all three enzymes on day 3 under BaP exposure, aligning with active degradation of the compound (Figure 1b). In contrast, activity levels remained unchanged in glucose-grown cells, and both gene expression and enzyme activity showed statistically significant differences, underscoring the specific effect of BaP. It is also worth noting that NADPH-cytochrome *c* reductase (CYP and CPR) and EH activities showed an early increase on day 1 in both conditions, which may reflect an early, non-specific cellular adjustment.

**Figure 2.**
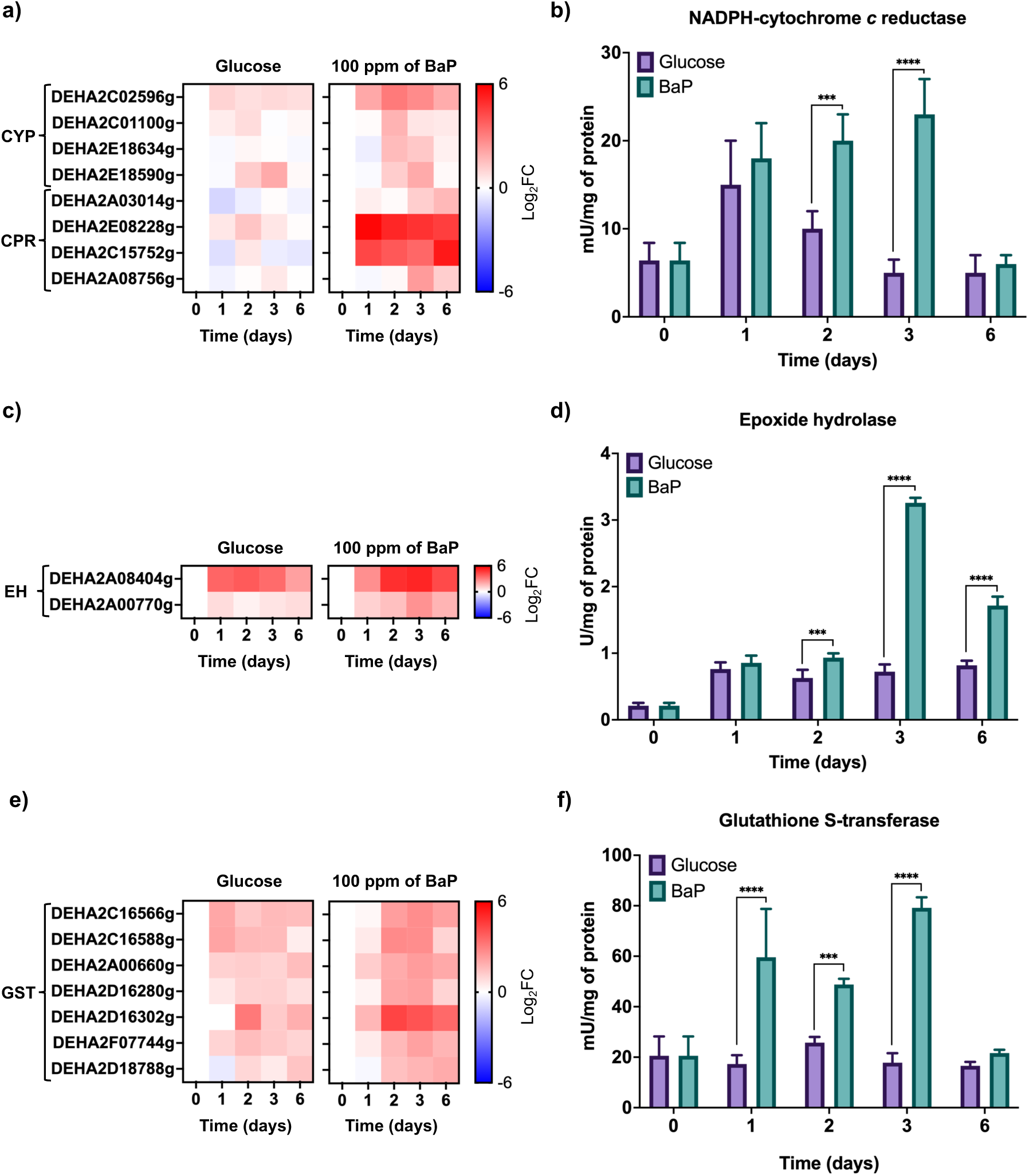
Differential gene expression and enzymatic activity of detoxification enzymes during BaP degradation in *D. hansenii*. a) Heatmap of cytochrome P450 (CYP) and cytochrome P450 reductase (CPR) gene expression under control (YNBG) and YNB + BaP (100 ppm)-treated conditions. b) NADPH-cytochrome c reductase activity measured over six days of growth. c) Heatmap of epoxide hydrolase (EH) gene expression under the same conditions. d) EH enzymatic activity over time. e) Heatmap of glutathione S-transferase (GST) gene expression. f) GST enzymatic activity profile. All RT-qPCR experiments were conducted with eight replicates (n = 8; four biological and two technical per condition), and expression values were normalized to *ACT1* (ORF DEHA2D05412g), control (YNBG) and YNB + 100 ppm BaP. Gene expression was quantified by RT-qPCR, normalized to *ACT1* (ORF DEHA2D05412g), using eight replicates (n = 8; four biological and two technical per condition). Heat maps were performed with data from Supplementary Figure 1. Enzymatic activities were assessed in three biological and three technical replicates (n = 9). Statistical analysis was performed using one-way ANOVA followed by Dunnett’s multiple comparisons test (*p* < 0.05). Asterisks indicate significance relative to control: (***) p = 0.0002; (****) p <0.0001.

GST plays a pivotal role in BaP detoxification by catalyzing the conjugation of GSH to its reactive metabolites (Capotorti et al. 2005). Exposure to BaP significantly increased the expression of GST-encoding ORFs (Figure 2e), which was accompanied by a corresponding rise in enzymatic activity (Figure 2f). These results pointed to a possible shift in glutathione homeostasis. To assess this, we quantified the levels of GSH and GSSG, calculated total glutathione (GSH + 2 × GSSG), and determined the GSH/GSSG ratio as an indicator of oxidative stress. In parallel, we evaluated the expression of the ORF encoding glutathione synthetase (GS) to determine whether *de novo* glutathione synthesis was activated (Figure 3).

**Figure 3.**
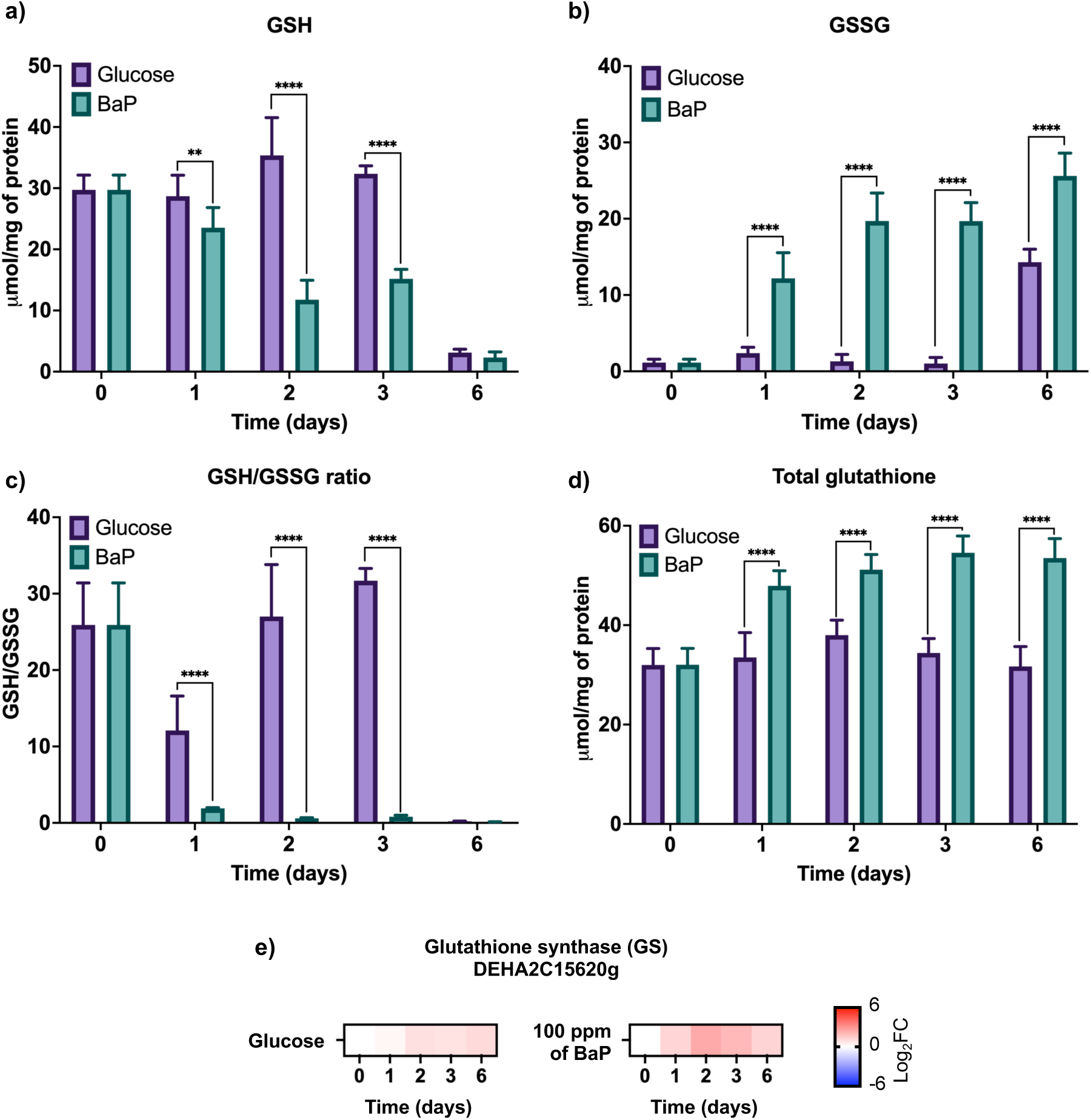
Changes in glutathione metabolism of *D. hansenii* in the presence of BaP. a) Reduced glutathione (GSH) levels along six days of growth in control (YNBG) and BaP-treated (YNB + 100 ppm BaP) conditions. b) Oxidized glutathione (GSSG) levels in the same conditions. c) GSH/GSSG ratio as a redox stress indicator, calculated from panels a) and b). d) Total glutathione content calculated as GSH + 2×GSSG. e) Relative expression of the glutathione synthetase (GS)-encoding ORF, measured by RT-qPCR. All biochemical measurements data were obtained from three biological and three technical replicates (n = 9). Gene expression was quantified by RT-qPCR, normalized to *ACT1* (ORF DEHA2D05412g), using eight replicates (n = 8; four biological and two technical per condition). The heat map was performed with data from Supplementary Figure 1. Statistical significance was determined by one-way ANOVA followed by Dunnett’s multiple comparisons test (*p* < 0.05). Asterisks indicate significance relative to control: (**) p = 0.0021; (****) p < 0.0001.

Under BaP exposure, we observed a significant decrease in GSH levels (Figure 3a). A notable decrease in GSH was observed, especially on day 6 in both conditions, which could reflect accumulated oxidative stress. A concomitant increase in GSSG, was also observed (Figure 3b), indicating active consumption of GSH to neutralize the contaminant. The GSH/GSSG ratio also dropped markedly (Figure 3c), revealing that *D. hansenii* was actively engaging its glutathione pool in detoxification and antioxidant defense mechanisms, hallmarks of a high oxidative stress environment. Interestingly, total glutathione content increased (Figure 3d), in parallel with the upregulation of GS expression (Figure 3e), suggesting enhanced glutathione biosynthesis as a compensatory response.

Given the observed disruptions in glutathione homeostasis, the next step was to examine the accumulation of ROS. BaP exposure led to a marked and statistically significant increase in both cytoplasmic (Figure 4a) and mitochondrial ROS (Figure 4b), as their different localization can reveal distinct sources and consequences of oxidative stress, detectable as early as day 1, before gradually declining. As elevated ROS levels can damage biomolecules, we quantified lipid peroxidation (Figure 4c) and carbonylated proteins (Figure 4d) as indicators of cellular damage. Both markers were elevated under BaP treatment, indicating that ROS accumulation was compromising membrane integrity and structural proteins.

**Figure 4.**
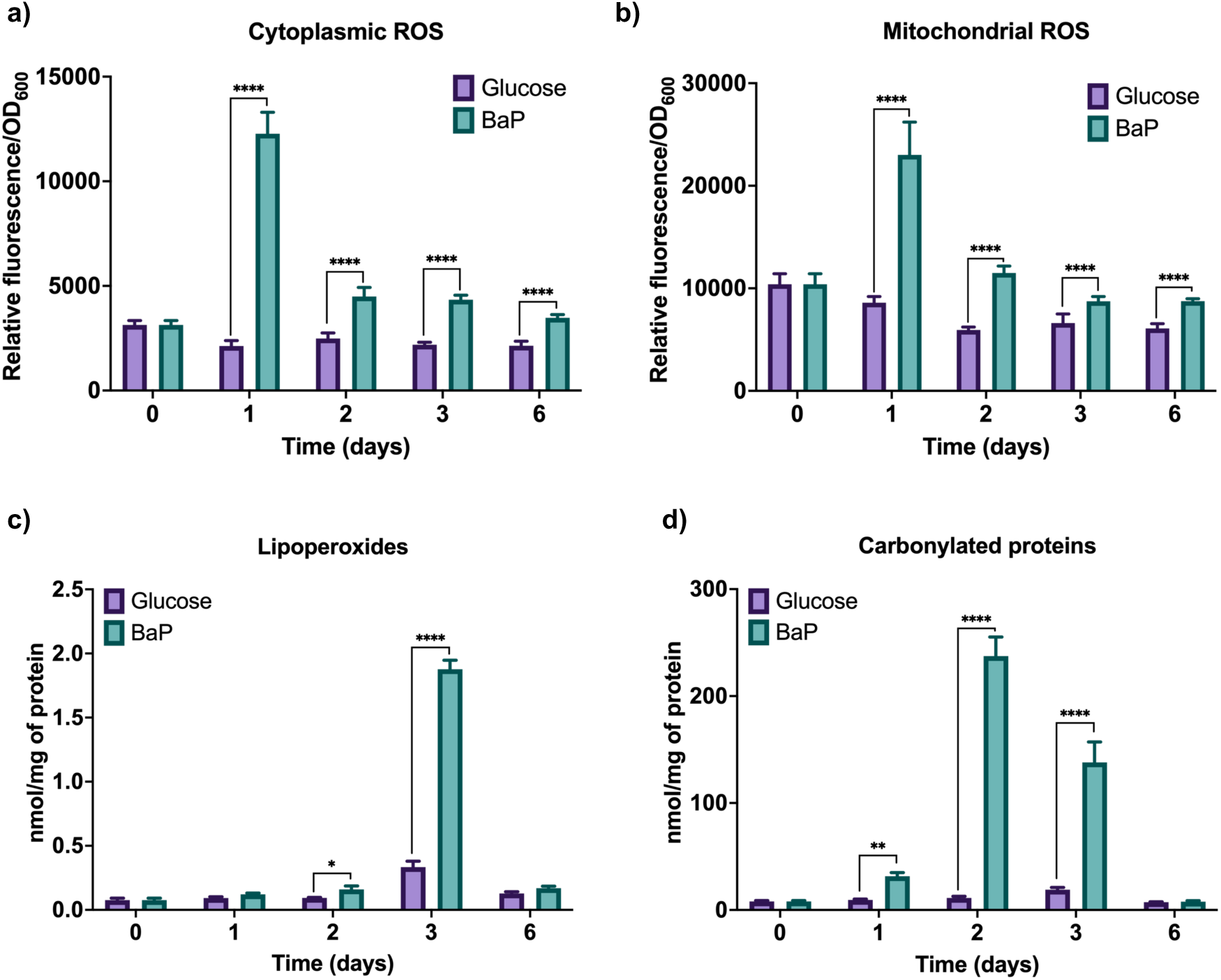
BaP exposure induces oxidative stress and molecular damage in *D. hansenii*. a) Cytoplasmic reactive oxygen species (ROS) levels detected using the H₂DCF-DA (2′,7′-dichlorodihydrofluorescein diacetate) probe during six days of growth in YNBG (control) and YNB + 100 ppm BaP. b) Mitochondrial ROS accumulation assessed using the DHR123 (Dihydrorhodamine 123) probe under the same conditions. c) Lipid peroxidation levels measured as the malondialdehyde (MDA) content, indicating membrane damage. d) Carbonylated proteins levels as a marker of oxidative protein damage. ROS levels were measured by fluorescence (H_2_DCF-DA: 435/530 nm; DHR123: 500/536 nm) and normalized to cell density (OD_600_). MDA and carbonylated proteins content were quantified spectrophotometrically (532 nm and 370 nm, respectively) and normalized to total protein. All experiments were performed using three biological and three technical replicates (n = 9). Statistical significance was determined using one-way ANOVA followed by Dunnett’s multiple comparisons test (p < 0.05). Asterisks indicate significance relative to control: (*) p = 0.0332; (**) p = 0.0021; (****) p < 0.0001.

The changes in glutathione homeostasis and ROS accumulation prompted us to examine the role of GR, the enzyme responsible for regenerating GSH from GSSG. A significant increase in GR transcript levels was observed on day 2 (Figure 5a), accompanied by a strong rise of enzymatic activity (Figure 5b). In parallel, GPx, which detoxifies organic peroxides, showed early transcriptional activation beginning on day 1 (Figure 5c), along with an increased enzymatic activity (Figure 5d). This response coincided with elevated lipid peroxidation, underscoring the intense oxidative pressure experienced by the cells during BaP metabolism.

**Figure 5.**
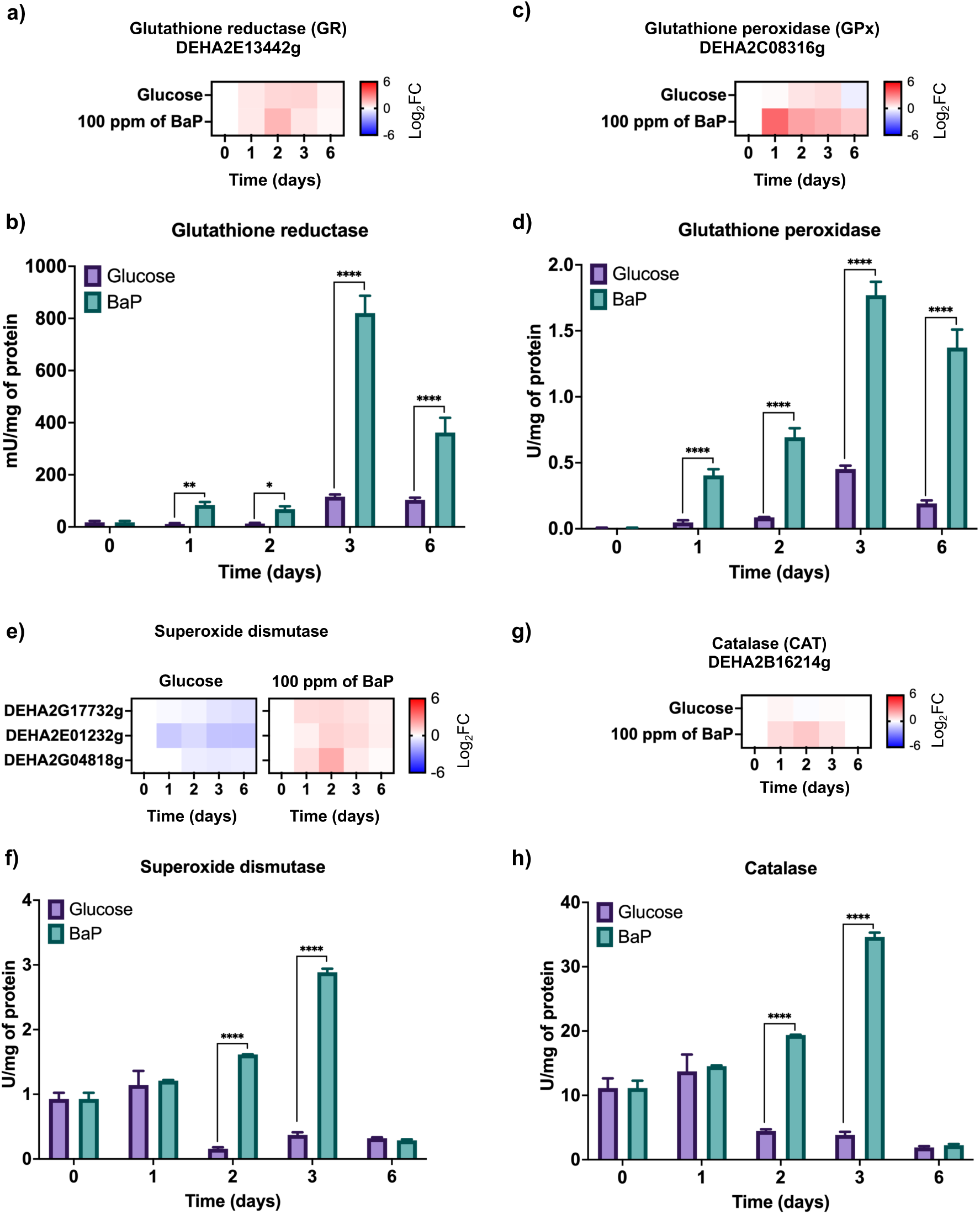
Activation of antioxidant defense enzymes in response to BaP-induced oxidative stress in *D. hansenii*. a) Relative expression of glutathione reductase (GR) measured by RT-qPCR. b) GR enzymatic activity across the six-day time course. c) Glutathione peroxidase (GPx) gene expression profile. d) GPx enzymatic activity e) Expression of superoxide dismutase (SOD)-encoding ORFs. (f) SOD enzymatic activity. g) Catalase (CAT) gene expression profile under control and BaP conditions. h) Catalase enzymatic activity. RT-qPCR was performed with eight replicates (n = 8; four biological and two technical per condition), using *ACT1* (ORF DEHA2D05412g) for normalization. Heat maps were performed with data from Supplementary Figure 1. Enzymatic activity assays were conducted using three biological and three technical replicates (n = 9). Statistical significance was assessed by one-way ANOVA followed by Dunnett’s multiple comparisons test (p < 0.05). Asterisks indicate differences relative to control: (*) p = 0.0332; (**) p = 0.0021; (****) p < 0.0001.

Changes in the expression and activity levels of SOD, where also observed. SOD is a key enzyme responsible for converting superoxide radicals (O_2_•⁻) into H_2_O_2_. The ORFs encoding SOD were upregulated under BaP exposure (Figure 5e), accompanied by a significant increase in enzymatic activity (Figure 5f). Similarly, CAT, which further breaks down H_2_O_2_ into H_2_O and O_2_, showed increased expression (Figure 5g) and activity (Figure 5h). These responses appear to be a direct consequence of ROS accumulation and BaP-induced molecular damage, highlighting the role of these enzymatic systems in protecting cells against oxidative injury.

Altogether, these findings suggest that BaP degradation in *D. hansenii* is not a passive metabolic event but rather a coordinated stress response focused on preserving cellular integrity. This response involves the transcriptional activation of multiple genes encoding enzymes such as CYP, CPR, EH, GST, GS, GR, GPx, SOD, and CAT. These proteins function collectively to facilitate xenobiotic detoxification, maintain glutathione homeostasis, and activate antioxidant defenses, thereby enhancing the yeast’s capacity to manage oxidative stress and metabolize BaP effectively. While these protective mechanisms support efficient detoxification and ROS neutralization, elevated levels of lipoperoxides and carbonylated proteins indicate that cellular damage still occurs. Together, these interconnected responses are essential for the survival and adaptation of *D. hansenii* in BaP-contaminated environments.

## 4. Discussion

Unlike previous studies, which focused on either gene identification or degradation rates, our work examines a broader approach by examining how *D. hansenii* responds to BaP at both the transcriptional and enzymatic levels. Notably, there is a clear coordination between detoxification pathways and antioxidant responses, suggesting a tightly linked cellular strategy that helps the yeast adapting to PAH stress. We focused on how *D. hansenii* responds to BaP, a toxic and persistent compound in both land and aquatic systems. (Cerniglia and Crow 1981; Mandal and Das 2018; Mandal et al. 2018; Padilla-Garfias et al. 2022; Padilla-Garfias et al. 2025).

Our results show that *D. hansenii* can tolerate 100 ppm of BaP without losing viability and can degrade nearly 70% of the compound within six days, even under glucose-limited conditions. This level of degradation is consistent with, and only slightly below the previously reported reduction of 84% achieved over ten days. This demonstrates the remarkable metabolic efficiency of this yeast. Interestingly, in previous work, *D. hansenii* demonstrated higher BaP degradation than several other yeast species, including *C. albicans* (77%), *R. mucilaginosa* (70%), and *S. cerevisiae* (79%) (Padilla-Garfias et al. 2022).

This tolerance appears to result from a coordinated metabolic adaptation, including the induction of key detoxification enzymes, such as CYP, CPR, EH, and GST, as well as a potent antioxidant defense system comprising of GR, GPx, SOD, and CAT. These enzyme systems act together with the critical antioxidant tripeptide glutathione, which plays a central role in maintaining intracellular redox balance (Schmacht et al. 2017; Padilla-Garfias et al. 2024). Not less important is that *D. hansenii* is a promising candidate for marine mycoremediation of PAH-contaminated environments because it can tolerate extreme conditions, including high salinity, oxidative stress, and nutrient limitation (Cerniglia and Crow 1981; Prista and Loureiro-Dias 2007; Segal-Kischinevzky et al. 2011; Bonugli-Santos et al. 2015; Sánchez et al. 2020; Martínez-Ávila et al. 2021; Padilla-Garfias et al. 2022; de la Fuente-Colmenares et al. 2024).

The ability of *D. hansenii* to grow in the presence of 100 ppm BaP (on both liquid and solid media) without significant loss of viability provides a strong physiological marker of tolerance to toxic compounds (Cerniglia and Crow 1981; Mandal and Das 2018; Mandal et al. 2018; Padilla-Garfias et al. 2022; Padilla-Garfias et al. 2025). Interestingly, when BaP was the only carbon source available, the yeast grew as much as it did with glucose as early as day two. This suggests that *D. hansenii* can not only tolerate BaP but also use it as a source of energy. This is like what has been seen in other fungi like *Aspergillus* sp. and *R. mucilaginosa* (Ostrem Loss et al. 2019; Martínez-Ávila et al. 2021; Peidro-Guzmán et al. 2021;

Padilla-Garfias et al. 2022; Padilla-Garfias et al. 2025). Similar PAH biodegradation capacity has been reported across several yeast genera, including *Candida guilliermondii*, *Yarrowia lipolytica* (formerly *Candida lipolytica*) (van der Walt and von Arx 1980; Zinjarde et al. 2014), *Candida maltosa*, *Candida tropicalis*, *Rhodotorula* spp., *Cryptococcus* spp., and *Debaryomyces* sp. itself, all of which have demonstrated efficient degradation of high-molecular-weight PAHs (Padilla-Garfias et al. 2024). This diversity of yeasts highlights the emerging importance of extremotolerant and extremophiles yeasts like *D. hansenii* in environments contaminated with PAHs, especially in marine or nutrient-limited environments.

The growth curve showed a significant increase in doubling time (from 7.5 hours in glucose to 18.3 hours in the presence of BaP). This suggests that the *D. hansenii* is adapting to the chemical stress by slowing down its metabolic processes, which is like what was seen under conditions of osmotic shifts and NaCl stress (Aggarwal and Mondal 2009; Sánchez et al. 2020). Throughout the six-day exposure to BaP, the total amount of biomass (as measured by dry weight) decreased, but cell growth continued steadily. This suggests that *D. hansenii* used its metabolic energy to support important processes like detoxification and survival, rather than using it to grow quickly (Ostrem Loss et al. 2019; Padilla-Garfias et al. 2022; Xiao et al. 2024). Fungi from the Ascomycota and Basidiomycota phyla exhibit similar growth patterns, with some species capable of surviving for over 40 days in the presence of anthracene (Godoy et al. 2016). Our findings suggest that *D. hansenii* has a flexible and stress-responsive growth strategy, and this strategy helps it to survive in toxic conditions by adjusting its physiological processes.

In Ascomycete fungi such as *D. hansenii*, the degradation of BaP and other xenobiotics has been linked to the activation of enzymatic pathways involved in sequential biotransformation processes (Morel et al. 2009; Morel et al. 2013; Aranda 2016; Padilla-Garfias et al. 2024). The results of this study show that the expression of ORFs related to detoxification enzymes, including CYP, CPR, EH, and GST, increased significantly starting on the third day of BaP exposure. This timing corresponds with the highest level of degradation seen in this and earlier studies (around 70%) (Padilla-Garfias et al. 2022; Padilla-Garfias et al. 2025). Gene expression, enzyme activity, and degradation kinetics all overlap, which supports the idea of a three-phase detoxification process. Phase I involves the oxidation of aromatic rings; phase II encompasses the hydrolysis of epoxides; and phase III is the conjugation of glutathione. Contrary to the notion of isolated events, these phases appear to develop in a coordinated manner (Morel et al. 2013; Aranda 2016; Padilla-Garfias et al. 2024). Although an increase in the enzymatic activities of NADPH-cytochrome *c* reductase (CYP and CPR) and EH was observed in both conditions, it should also be noted that these enzymes participate in secondary metabolism, which may partially explain their basal activity in the control medium (Fretland and Omiecinski 2000; Durairaj et al. 2016).

Interestingly, similar timing patterns have been reported in other fungi such as *Aspergillus* species, *R. mucilaginosa*, and even in previous work with *D. hansenii*. In all cases, the peak expression of CYP genes tends to coincide with the highest rates of PAH degradation, suggesting that this phenomenon is a conserved adaptive feature among different fungal species (Capotorti et al. 2005; Ostrem Loss et al. 2019; González et al. 2020; Martínez-Ávila et al. 2021; González et al. 2021; Peidro-Guzmán et al. 2021; Padilla-Garfias et al. 2022; González et al. 2022). In contrast, control cells that were cultured in the absence of BaP exhibited minimal alterations in gene expression or enzymatic activity. This finding corroborates the notion that the observed responses are not constitutive but rather are specifically triggered by the presence of the contaminant (Miglani et al. 2022).

Within the group of enzymes activated by BaP exposure, the GSTs were the most important enzymes that were affected by BaP exposure. Their increased activity, both at the gene expression level and in enzymatic activity, suggests that they play a key role in detoxifying BaP by attaching glutathione to its reactive byproducts. This step is essential for neutralizing those intermediates and directing them either toward export, sequestration in the vacuole, or further metabolic breakdown (Zablotowicz et al. 1995; Capotorti et al. 2005; Allocati et al. 2018; Mazari et al. 2023; Padilla-Garfias et al. 2024).

While detailed functional analysis of each regulated ORF is beyond the scope of this study, our results highlight the coordinated transcriptional activation of multiple detoxification- and antioxidant-related genes, supporting a systems-level adaptation strategy.

As GST expression and activity ramped up in response to BaP, we also saw a clear shift in the glutathione balance inside the cell. GSH levels dropped significantly, while GSSG levels rose, leading to a lower GSH/GSSG ratio, a classic indicator that the cell is under oxidative stress. The significant drop in GSH levels by day 6 under both BaP and glucose conditions likely reflects a sustained oxidative challenge that progressively overwhelms the yeast’s antioxidant defenses. (Livingstone 2001; Li et al. 2009; Breitenbach et al. 2015; González et al. 2020; Binati et al. 2022; Yaakoub et al. 2022). At the same time, we observed increased expression of GS, suggesting that the yeast was trying to restore its GSH pool through *de novo* synthesis; despite an initial compensatory increase in total glutathione and GS expression, prolonged exposure appears to exhaust the cellular glutathione pool, indicating a critical tipping point in redox homeostasis. Even though GS itself isn’t the rate-limiting enzyme in the glutathione pathway (Kurylenko et al. 2019; Wangsanut and Pongpom 2022).

The presence of BaP resulted in a substantial increase in ROS within *D. hansenii*, as evidenced by their detection in both the cytosol and mitochondria. This observation indicates a pervasive oxidative stress within the cell (Jamieson 1998; Herrero et al. 2008; Aranda 2016; Yaakoub et al. 2022; Padilla-Garfias et al. 2024; Padilla-Garfias et al. 2025). Interestingly, ROS levels increased early, peaking on day one and then decreasing slightly by day two. But even after that, they remained higher than in untreated cells throughout the rest of the experiment. Measuring both cytoplasmic and mitochondrial ROS provided insight into the compartment-specific origins and persistence of oxidative damage (Yaakoub et al. 2022). This prolonged oxidative pressure started to take a visible toll; membrane integrity and protein structure were clearly affected. We saw strong increases in lipid peroxidation, as indicated by MDA levels, and in protein carbonylation, both classic signs of oxidative stress and cellular damage (Jamieson 1998; Salmon et al. 2004; Ikner and Shiozaki 2005; Herrero et al. 2008).

The initial accumulation of ROS and the concomitant oxidative damage to essential biomolecules provide substantial evidence that BaP metabolism triggers oxidative stress, presumably resulting from the production of reactive intermediates during CYP-mediated phase I oxidation. Similar oxidative responses have been documented in other eukaryotic models, including *S. cerevisiae*, *Dentipellis* sp., *Aspergillus* sp., and *R. mucilaginosa*, where BaP or PAH exposure leads to ROS accumulation and activation of antioxidant pathways (Park et al. 2019). Interestingly, even the green alga *Ulva lactuca* shows a comparable redox imbalance under PAH stress, suggesting that this type of oxidative response might be conserved across quite distant lineages, from fungi to algae (González et al. 2020; González et al. 2021; González et al. 2022).

Although a significant increase in ROS was observed in *D. hansenii* treated with BaP compared to the control group, a reduction in ROS levels was observed cells treated with BaP after day 2. This indicates the effective and timely activation of antioxidant defense mechanisms to prevent cellular damage and restoring intracellular redox balance (Jamieson et al. 1996; Emri et al. 1997; Yaakoub et al. 2022). The antioxidant response in *D. hansenii* exhibited a sequential and coordinated pattern, with the expression and activity of GPx increasing as early as day 1. This was followed by the induction of GR, SOD, and CAT, which peaked between days 2 and 3. The observed decline in both cytoplasmic and mitochondrial ROS has also been shown to correlate with the upregulation and increased activity of compartment-specific antioxidant enzymes. The temporal cascade suggests a tiered oxidative defense strategy, fine-tuned to neutralize the rising ROS burden and the secondary oxidative byproducts generated during phase I and II detoxification processes (Herrero et al. 2008; Yaakoub et al. 2022).

The maintenance of redox balance in *D. hansenii* appears to be predominantly dependent on the glutathione system. The shifts in GSH and GSSG levels, coupled with the upregulation of GS and GR, indicate that this pathway functions in a dual capacity. On one hand, it has been demonstrated to facilitate the cell’s ability to manage oxidative stress, and on the other hand, it has been shown to ensure the continued operation of the detoxification mechanisms in the presence of BaP exposure. This phenomenon is not simply a protective mechanism; rather, it functions as a central coordinator that integrates redox regulation with xenobiotic processing. This integrative role of glutathione metabolism complements the phase I–III detoxification enzymes, reinforcing its place as a critical control point in the cell’s adaptive response (Emri et al. 1997; Kurylenko et al. 2019; Binati et al. 2022; Wangsanut and Pongpom 2022).

The observations made in *D. hansenii* are consistent with the responses exhibited by other extremotolerant yeasts, such as *Y. lipolytica* and *Debaryomyces fabryi*. These organisms are characterized by their robust antioxidant defenses and their capacity to metabolize xenobiotics in adverse environmental conditions. (Sekova et al. 2015; Yaakoub et al. 2022). This phenomenon of interspecies convergence, both within and across the *Debaryomyces* genus, serves to substantiate the notion that these responses are neither arbitrary nor isolated. Instead, these findings appear to reflect an evolutionarily conserved survival strategy among fungi that thrive in extreme or contaminated niches. In addition to fungi, this effect has also been observed in some plants, such as *Arabidopsis thaliana*, and algae, such as *U. lactua* (Skipsey et al. 2011; González et al. 2020; González et al. 2021; González et al. 2022).

Beyond its ability to degrade BaP and linear hydrocarbons and withstand considerable environmental stress, *D. hansenii* demonstrates clear advantages over other eukaryotic degraders, including *Aspergillus*, *Rhizopus*, and *Candida* species, thanks to its unique physiological resilience and metabolic versatility (Padilla-Garfias et al. 2024). Its marine provenance endows it with a resilience to osmotic and nutritional stress, a trait that is imperative for bioremediation in saline and nutrient-limited environments (Aggarwal and Mondal 2009; Sánchez et al. 2020). Despite the complexity of genetic manipulation, successful mutant strains have been generated (Sánchez et al. 2020), thereby establishing the foundation for future metabolic engineering endeavors. The enhancement of its biodegradation potential can be achieved through the optimization of culture conditions, microbial consortia, or synthetic biology approaches. To sum up, the characteristics serve to substantiate *D. hansenii* as a compelling contender for PAH bioremediation in ecologically demanding environments.

These results support an integrative model in which the “xenome” of *D. hansenii*, as well as its inducible genetic and enzymatic repertoire for xenobiotic processing, act as a dynamically regulated system in response to BaP exposure (Morel et al. 2013; Labrou et al. 2015; Aranda 2016; Padilla-Garfias et al. 2021; Padilla-Garfias et al. 2024). As illustrated in Figure 6, the detoxification cascade comprises three interconnected phases: (I) aromatic ring oxidation, (II) epoxide hydrolysis, and (III) glutathione conjugation. These metabolic pathways are intricately linked with antioxidant defenses, particularly the glutathione system and enzymes such as CYP, EH, GST, GR, GPx, SOD, and CAT.

**Figure 6.**
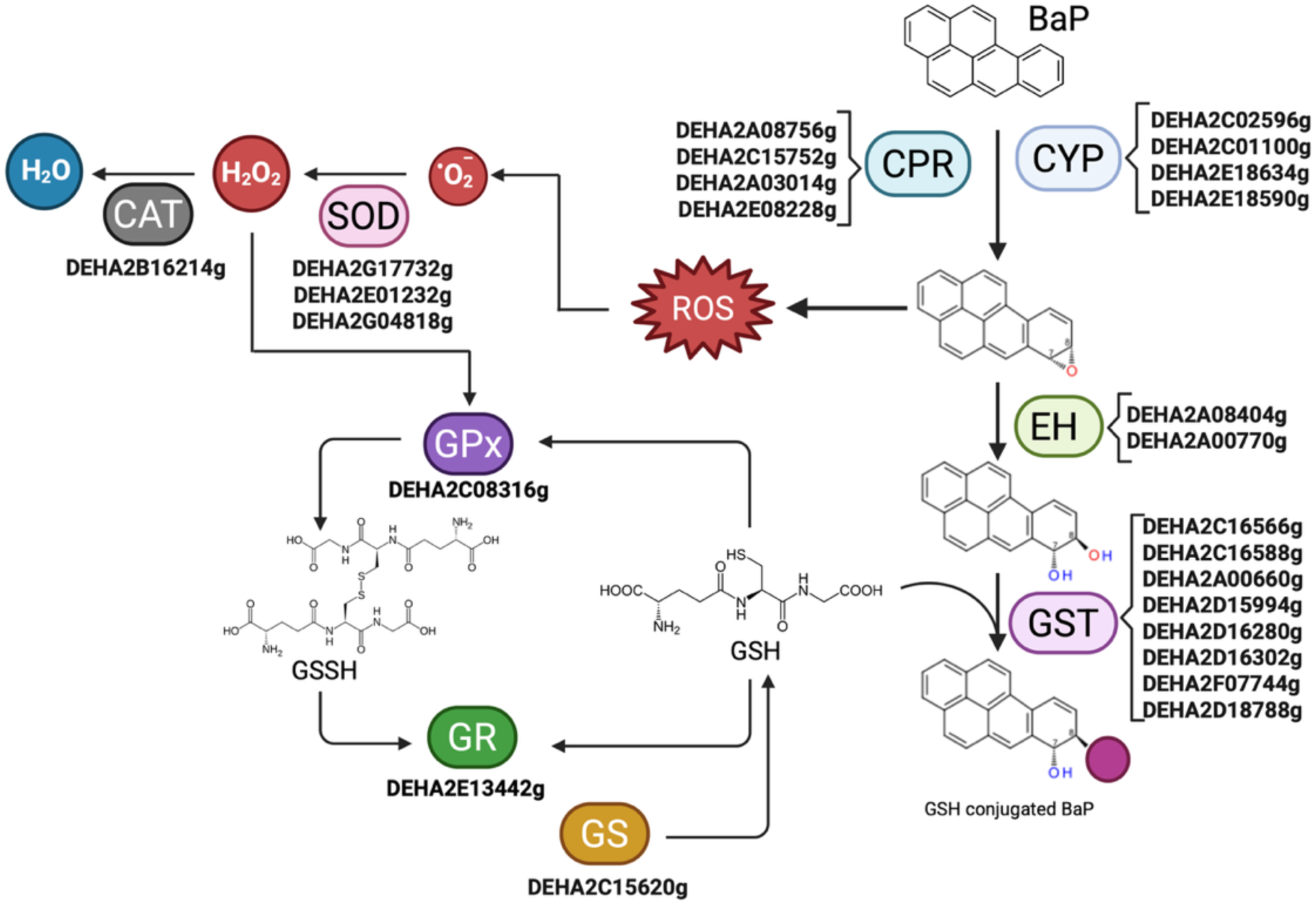
Schematic representation of the coordinated detoxification and antioxidant response of *D. hansenii* to BaP exposure. The model illustrates a three-phase metabolic pathway: (I) BaP oxidation by cytochrome P450 monooxygenases (CYPs), (II) hydrolysis of reactive epoxides by epoxide hydrolases (EHs), and (III) glutathione-mediated conjugation of BaP metabolites via glutathione S-transferases (GSTs) (the pink circle represents the GSH molecule). These detoxification routes are functionally coupled with the activation of antioxidant defenses, including glutathione peroxidase (GPx), glutathione reductase (GR), superoxide dismutase (SOD), and catalase (CAT), which collectively mitigate ROS accumulation and preserve redox homeostasis.

This coordination suggests that the effective BaP detoxification in *D. hansenii* is contingent not only on the enzymatic conversion of the xenobiotic but also on the preservation of cellular redox balance under oxidative challenge. The simultaneous modulation of detoxification and antioxidant gene networks unveils a sophisticated regulatory strategy that fosters cell survival in toxic environments. *D. hansenii*’s adaptive molecular profile, which is characterized by its extreme tolerance, positions it as a valuable eukaryotic model for studying xenobiotic metabolism. Furthermore, it is a promising candidate for biotechnological applications in marine mycoremediation (Aranda 2016; Padilla-Garfias et al. 2022; Navarrete et al. 2022; Padilla-Garfias et al. 2024; Padilla-Garfias et al. 2025).

## 5. Conclusions

*D. hansenii* not only tolerates the presence of BaP, but also actively degrades it by mounting a tightly regulated response that integrates enzymatic detoxification with antioxidant defense, even under carbon-starved conditions such as the absence of glucose. By combining molecular and biochemical analyses, we identified the involvement of CYP, CPR, EH, GST, glutathione metabolism, and key antioxidant enzymes. These elements coordinately operate to manage both the metabolic demands of BaP degradation and the oxidative stress it provokes.

Beyond isolated molecular observations, our findings converge into a conceptual model that illustrates how *D. hansenii* combines detoxification and redox regulation mechanisms to preserve cellular viability in chemically adverse environments. This integrated strategy highlights not only the remarkable adaptive capacity of this marine yeast but also reinforces its potential as a robust microbial candidate for mycoremediation applications.

## Acknowledgements

The authors gratefully acknowledge Dr. Diego González-Halphen, Dr. James González and Dr. Augusto César Poot-Hernández for their valuable scientific guidance. We also thank Dr. Laura Ongay-Larios, MSc. Minerva Mora, BSc. Guadalupe Codiz-Huerta, Dr. Carlos Alberto Peralta-Álvarez, Dr. Martha Lucinda Contreras-Zentella, Dr. Carlos Campero-Basaldua and Eduardo Cardona Guerrero for their technical assistance. The support of Dra. Claudia Segal-Kischinevzky and Dra. Alicia González Manjarrez with the performance of some the experiments in this work is also gratefully acknowledged.

This manuscript was written by the authors. Generative AI tools (ChatGPT-4 and DeepL Write) were only used to refine language and clarity, with all scientific content authored and reviewed by the authors. Additionally, Mendeley was employed for the management of bibliographic references, and BioRender.com was employed to create Figure 6.

## 6. Author Contributions

Conceptualization: F.P.-G., A.P.; Data curation: F.P.-G.; Formal analysis: F.P.-G.; Funding acquisition: A.P.; Investigation: F.P.-G., M.A.-V., M.C., N.S.S.; Methodology: F.P.-G.; Project administration: A.P.; Resources: A.P.; Supervision: A.P.; Validation: F.P.-G., M.A.-V., M.C., N.S.S., A.P.; Visualization: F.P.-G.; Writing – original draft: F.P.-G., M.C., A.P.; Writing – review & editing: F.P.-G., M.A.-V., M.C., N.S.S., A.P. All authors read and approved the final version of the manuscript.

## 7. Funding

This research was supported by grants IN204321 and IN217924 from the Dirección General de Asuntos del Personal Académico (DGAPA), Universidad Nacional Autónoma de México (UNAM). Francisco Padilla-Garfias is a doctoral student in the Programa de Maestría y Doctorado en Ciencias Bioquímicas at UNAM and receives a scholarship (CVU 904691) from the Secretaría de Ciencia, Humanidades, Tecnología e Innovación (SECIHTI). Minerva Georgina Araiza-Villanueva acknowledges postdoctoral support from DGAPA-UNAM.

## 8. Ethics Statement

All experimental procedures involving *Debaryomyces hansenii* were conducted in accordance with the institutional biosafety guidelines of the Instituto de Fisiología Celular, UNAM. The yeast strain used in this study is classified as non-pathogenic and did not require specific ethical approval under current national or international regulations.

## 9. Conflicts of Interest

The authors declare no conflicts of interest.

---

**Supplementary Table 1.**
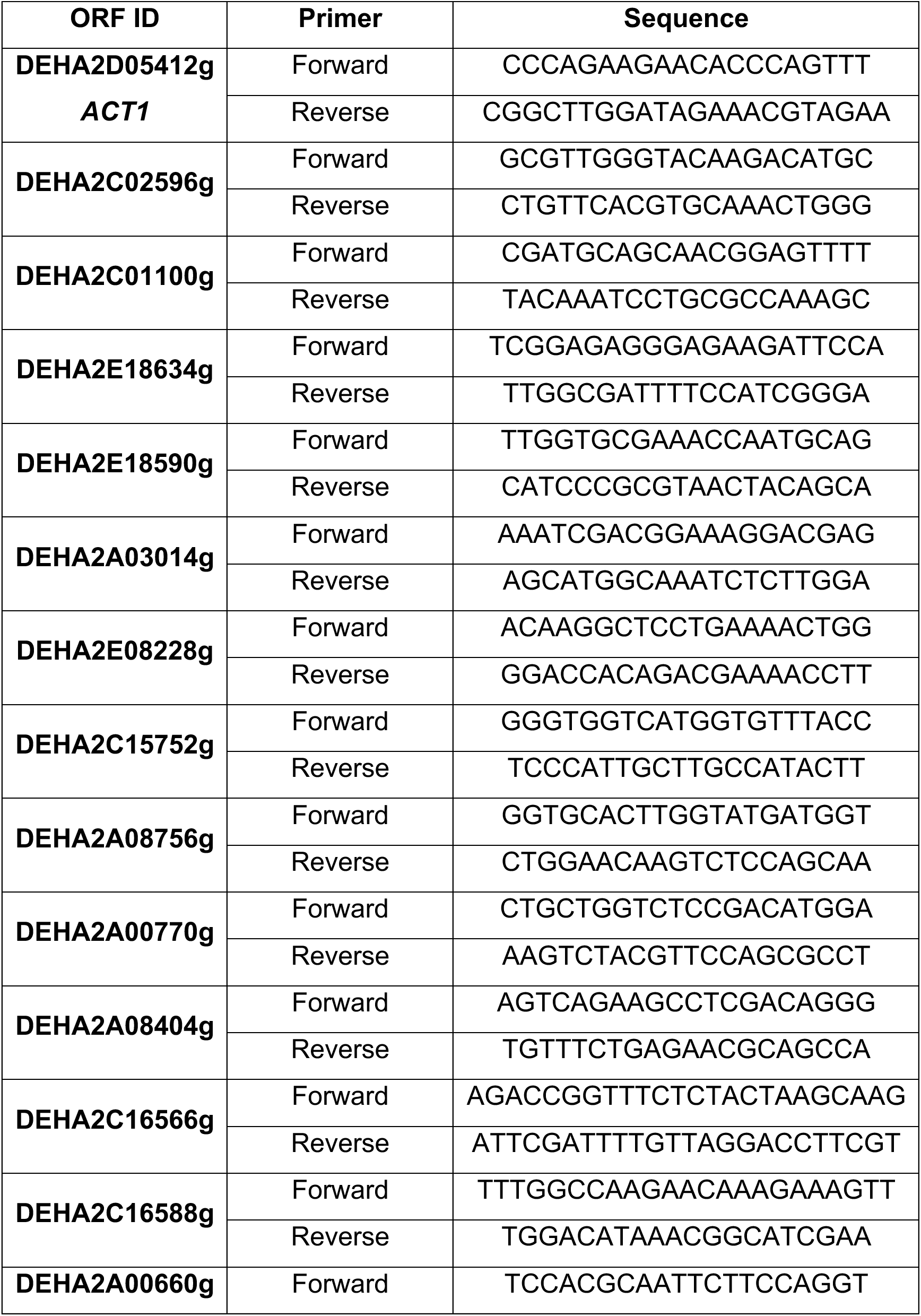
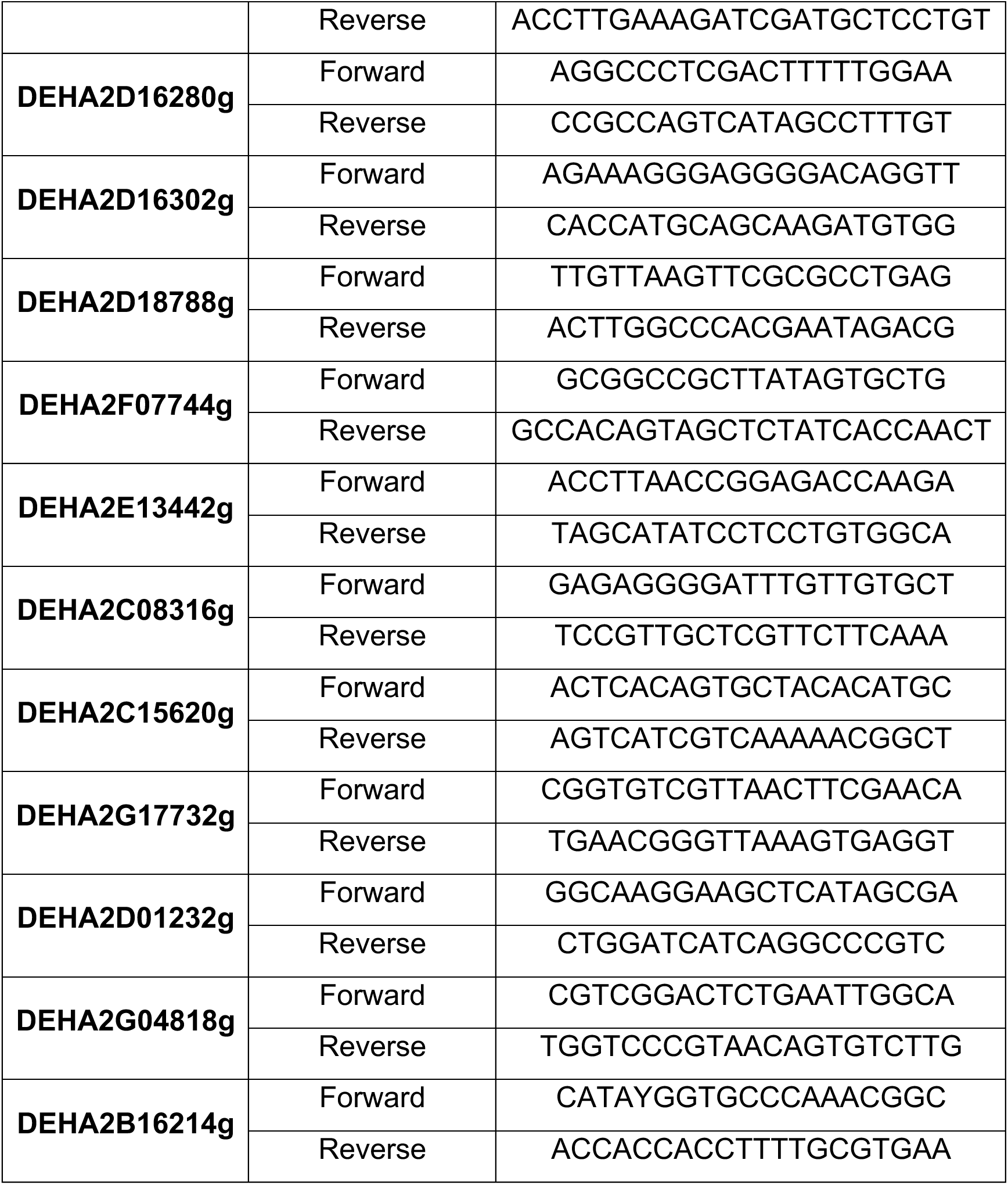
Oligonucleotides used in qPCRs.

**Supplementary Table 2.**
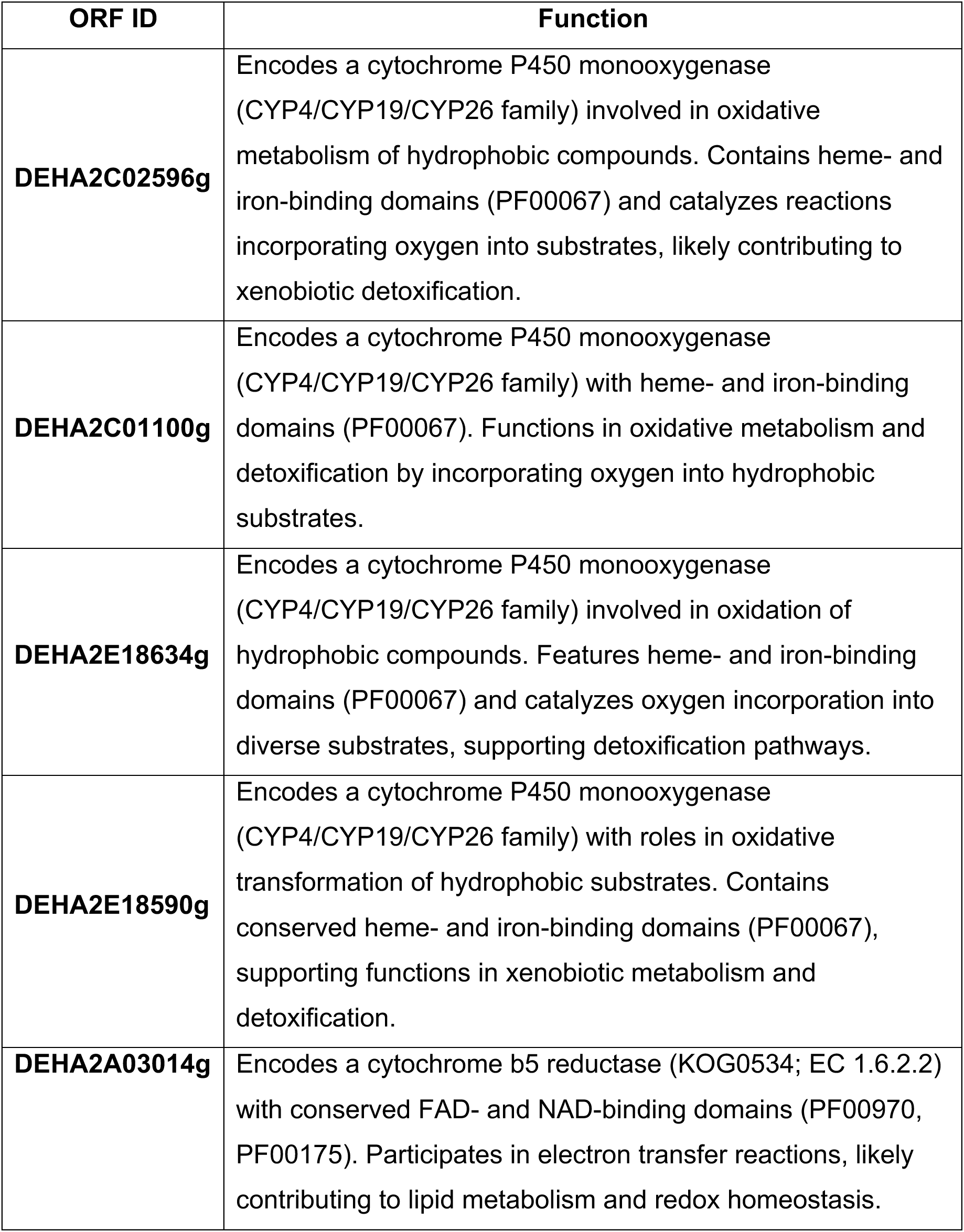
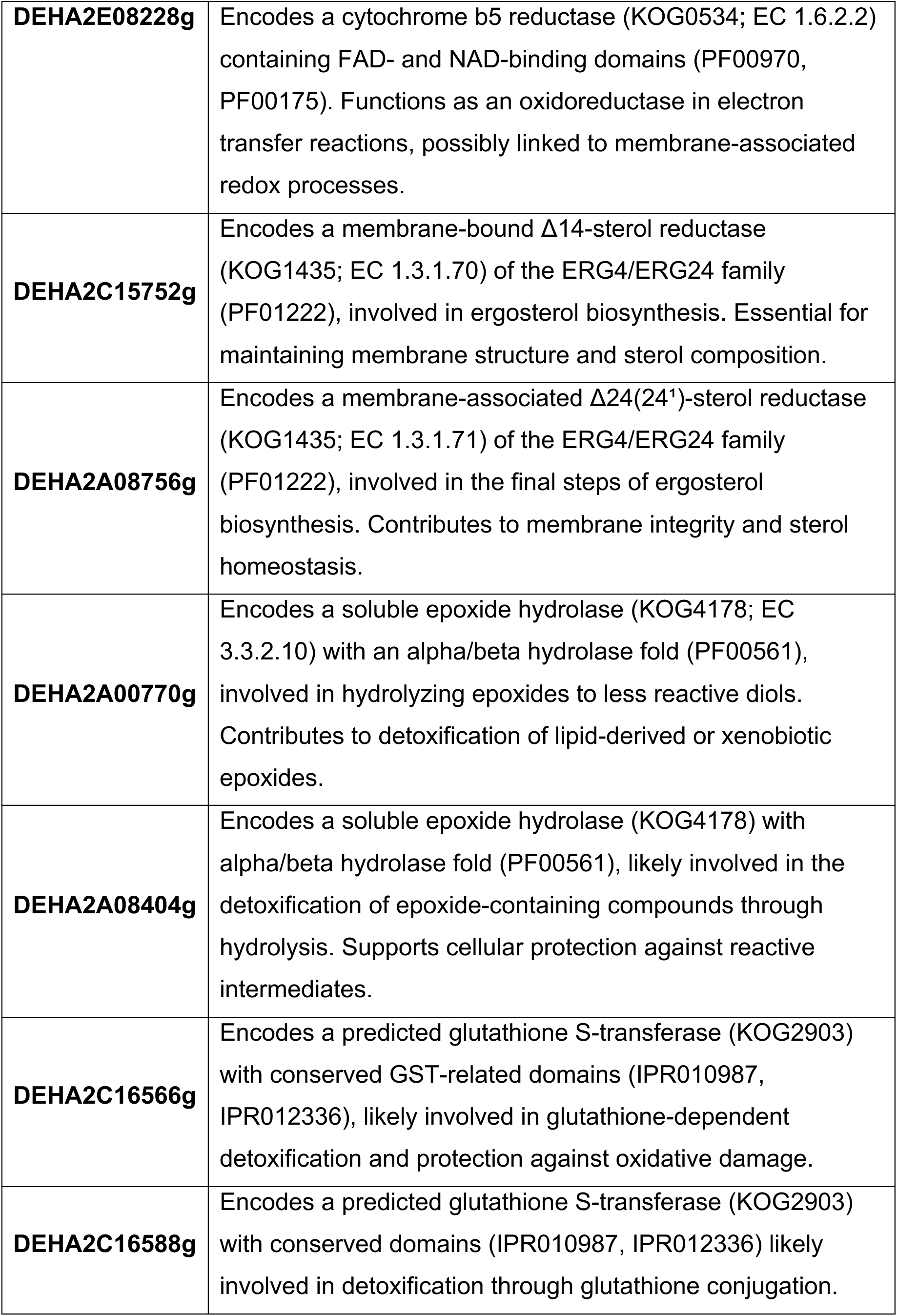
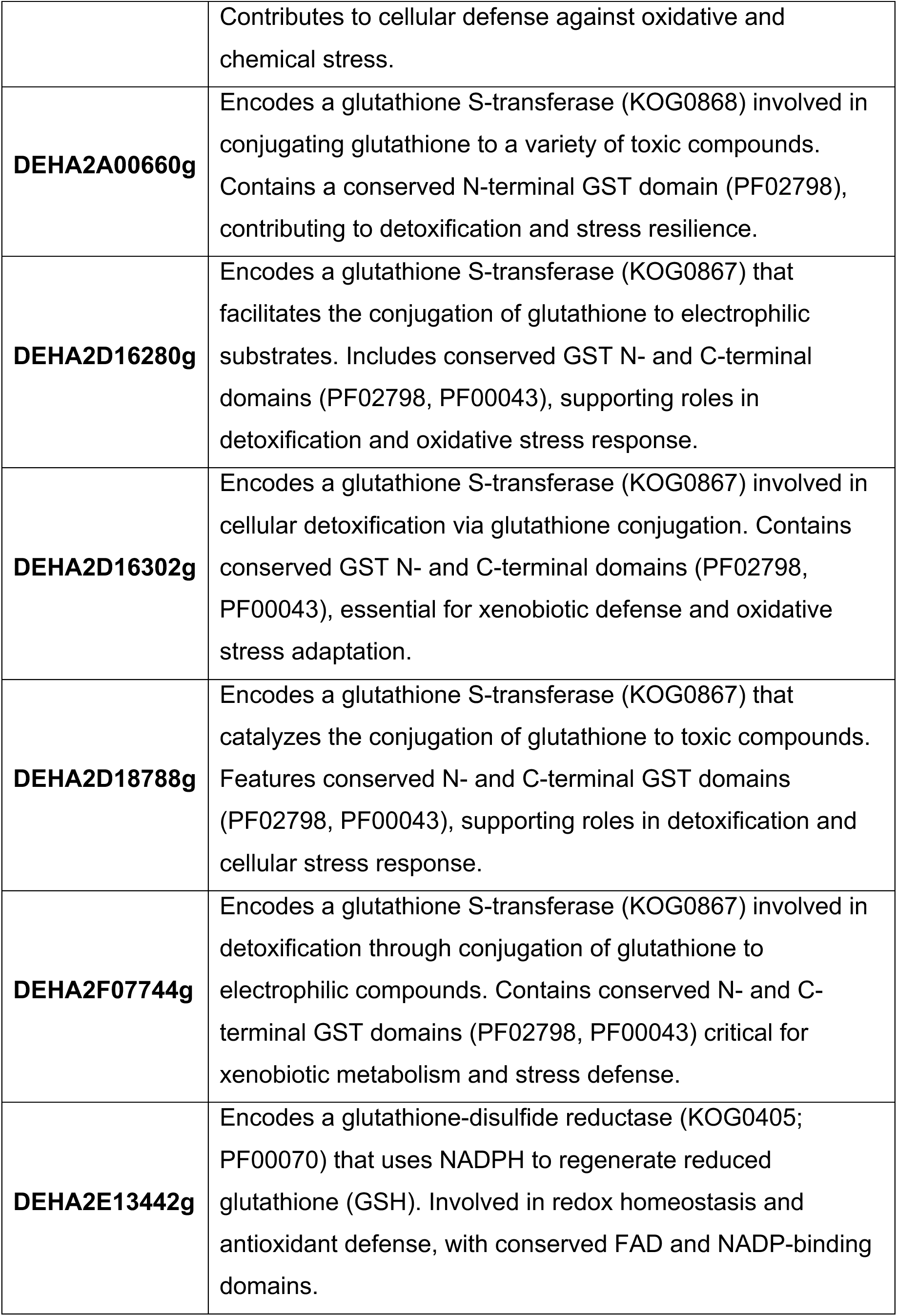
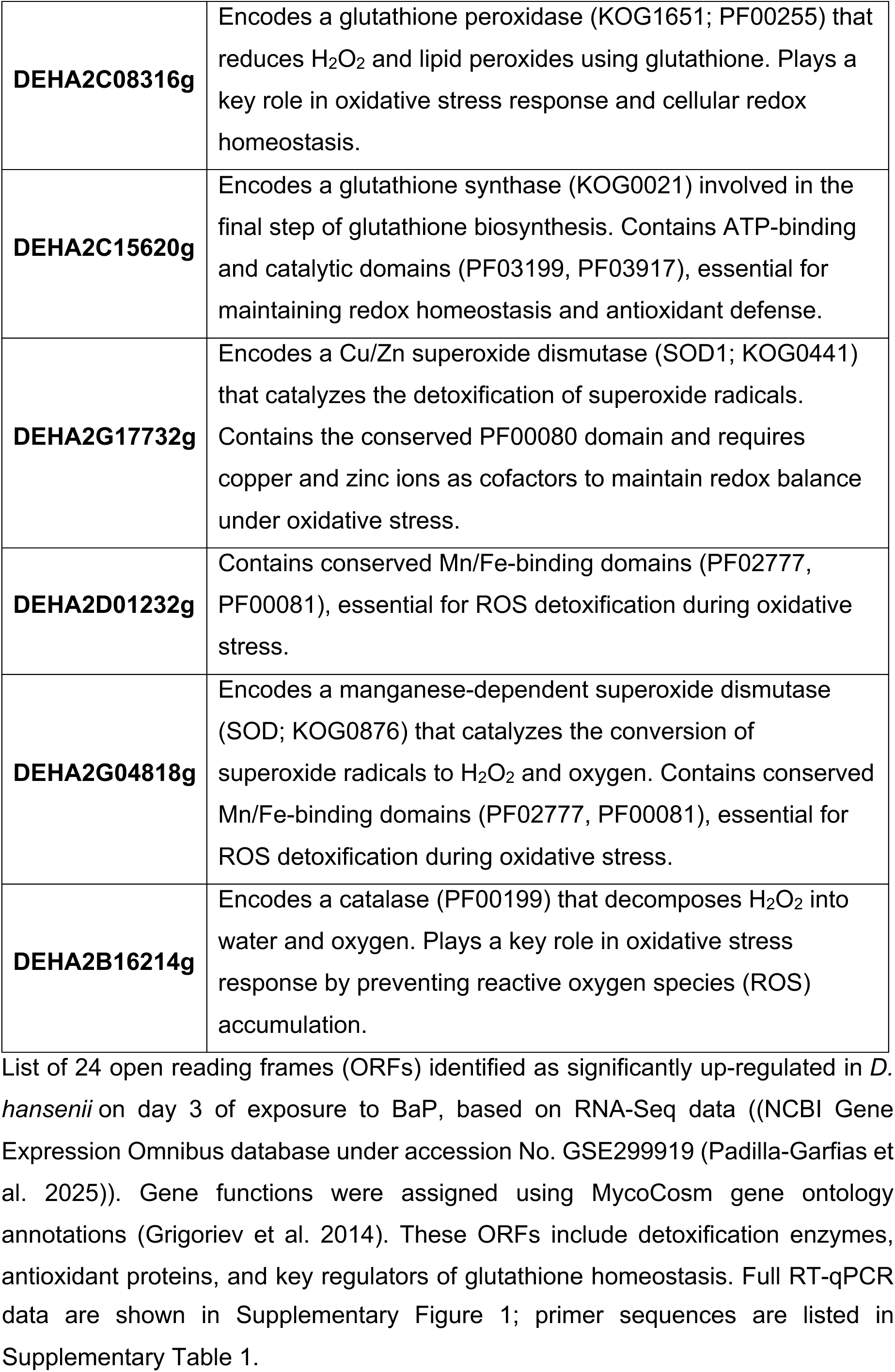
Function of 24 up-regulated genes in *D. hansenii* associated with BaP detoxification, glutathione metabolism, and antioxidant defense.

**Supplementary Figure 1.**
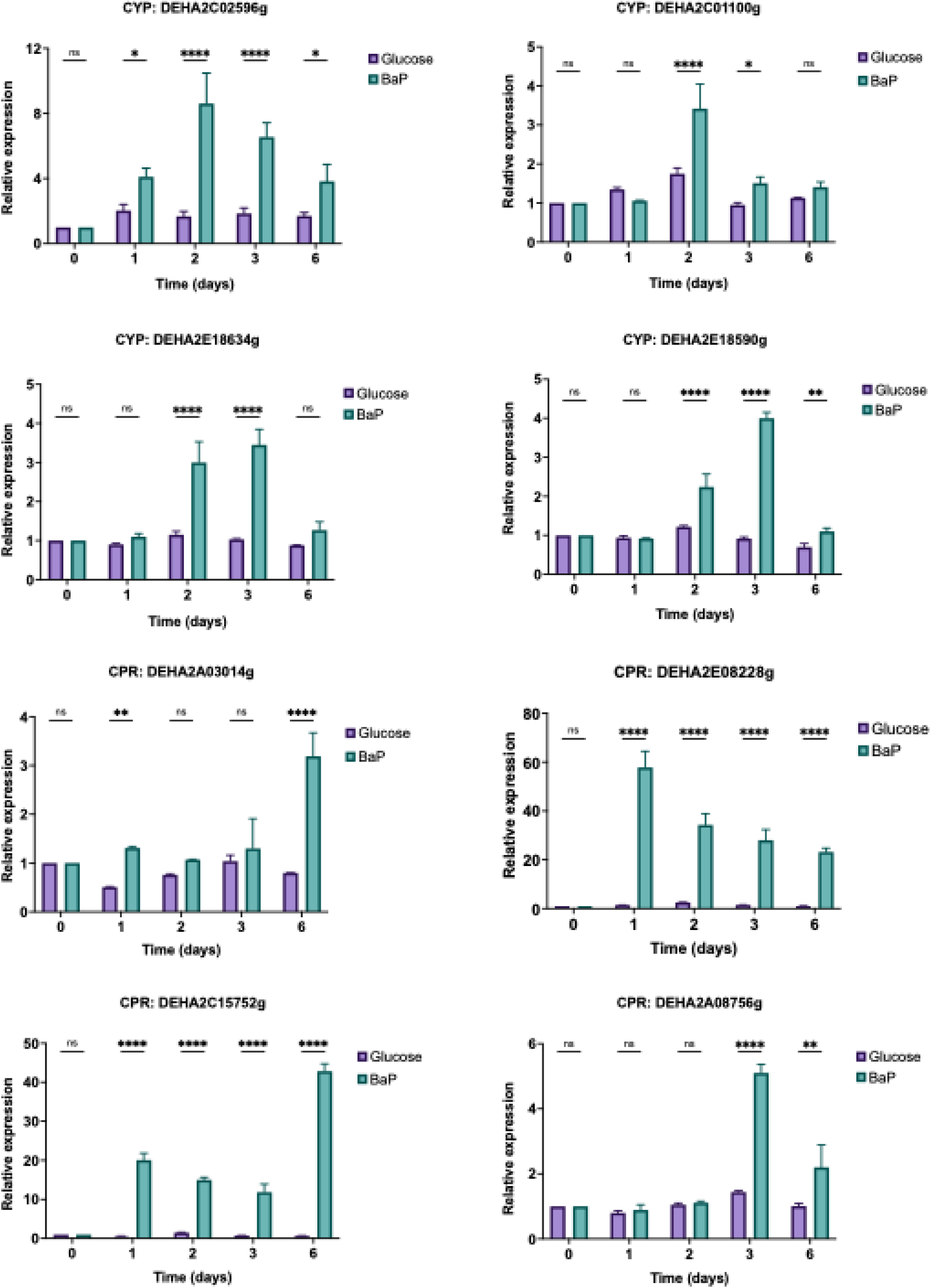
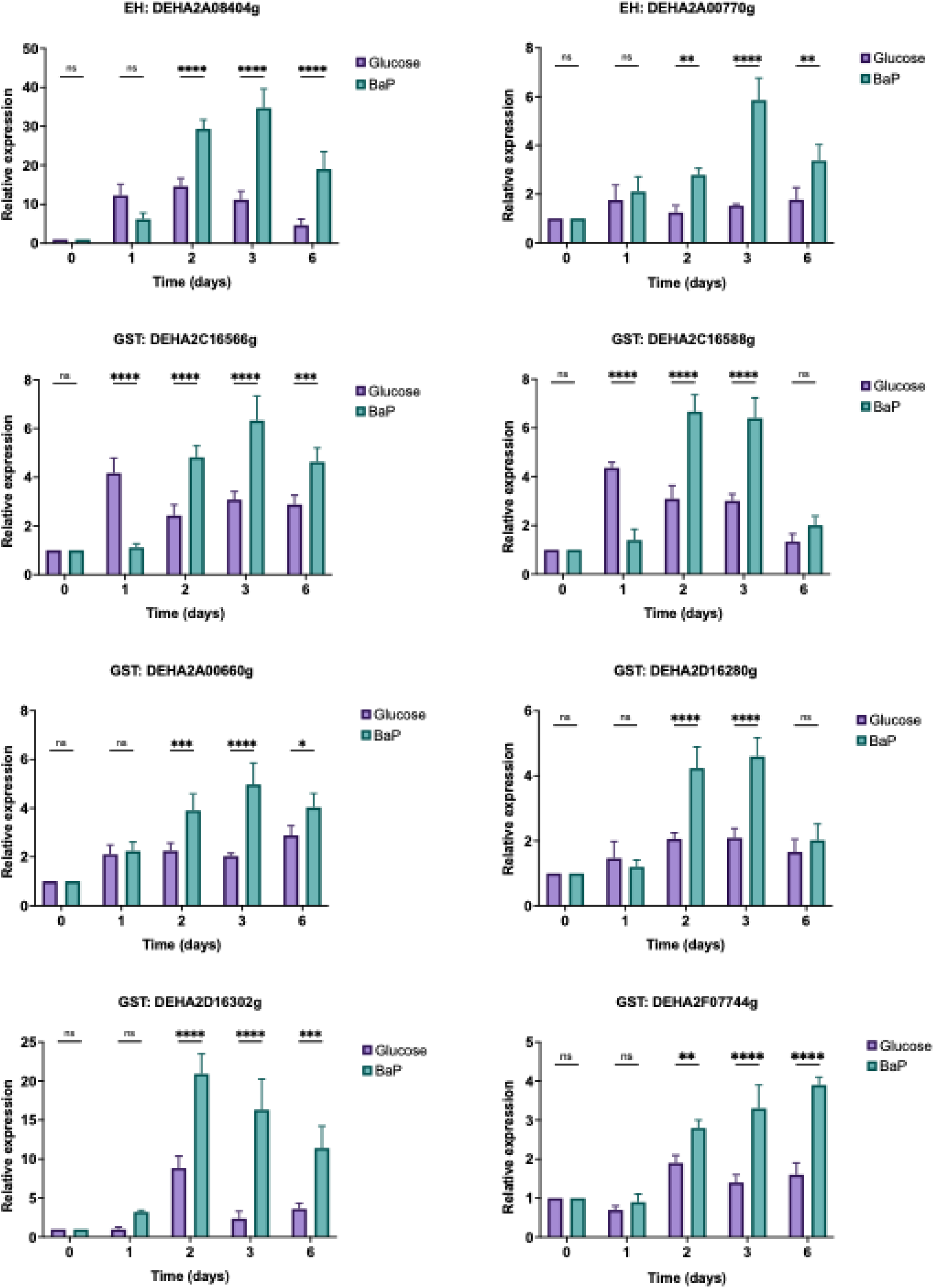
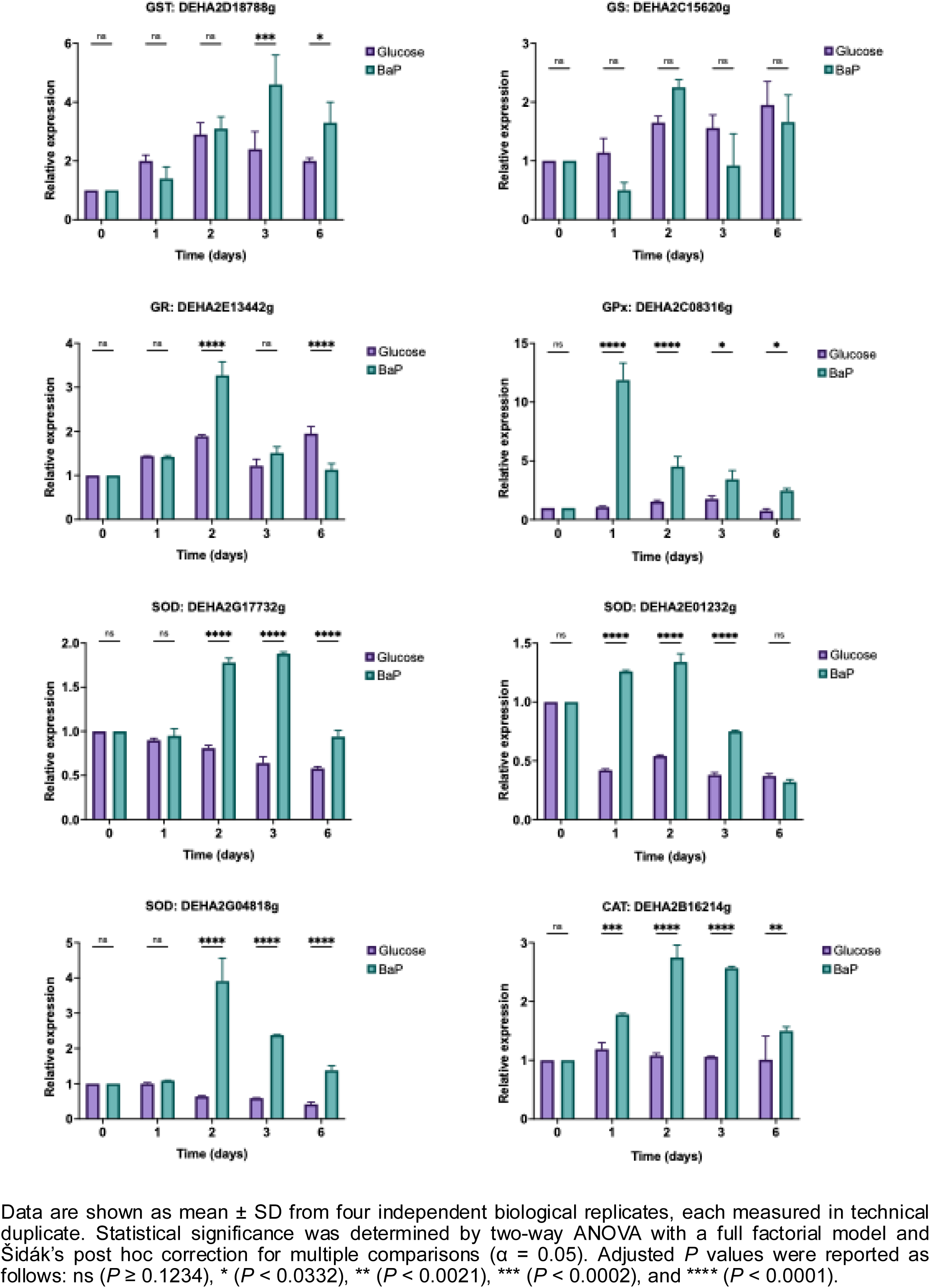
RT-qPCR data used to construct the heat maps shown in the article.

## Notes

### Competing Interest Statement

The authors have declared no competing interest.

