## Supplementary Figure 1 for "Benzo(a)pyrene degradation induces coordinated antioxidant and detoxification responses in the marine yeast *Debaryomyces hansenii*"

**Supplementary Figure 1.** RT-qPCR data used to construct the heat maps shown in the article.

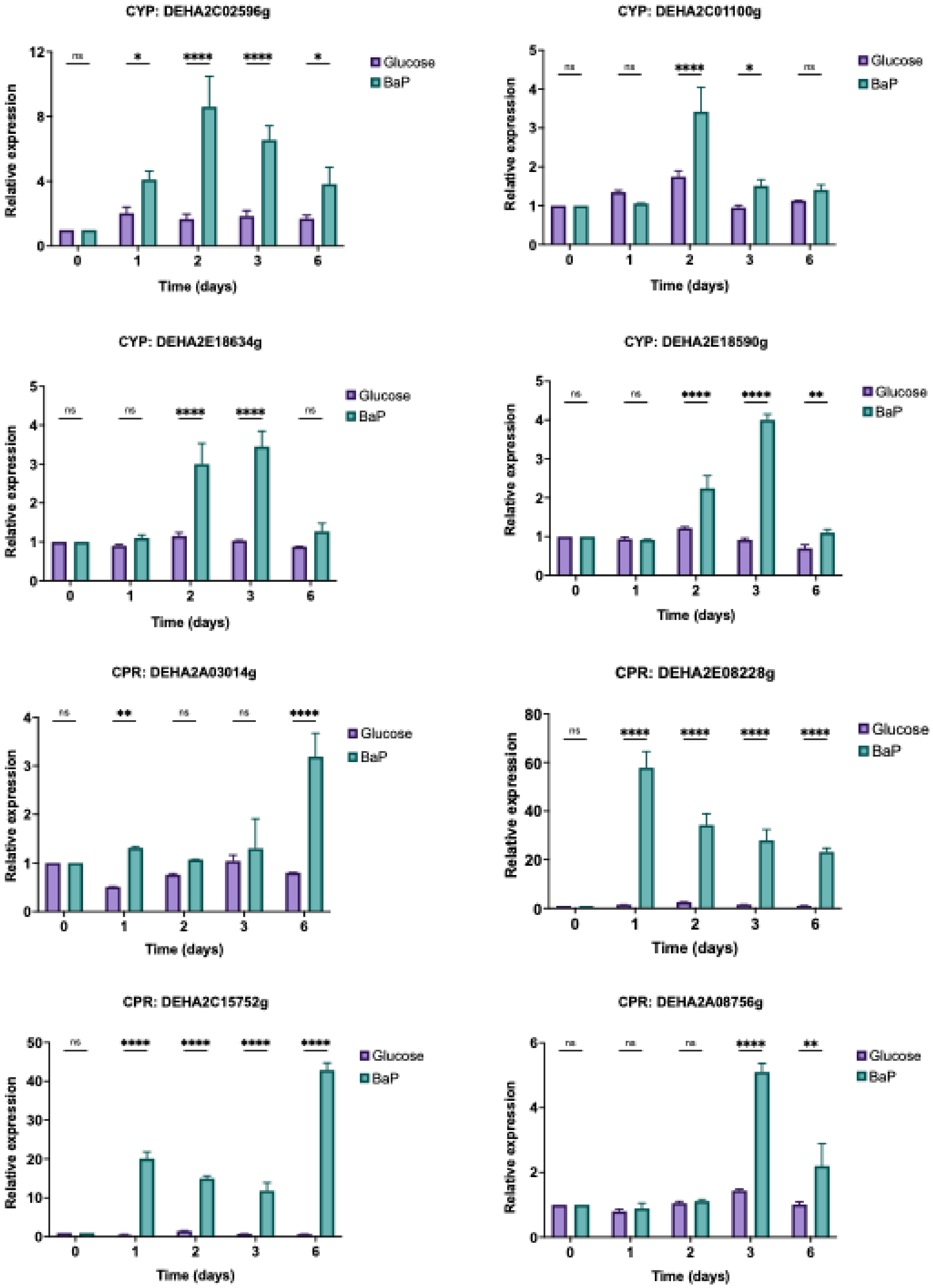

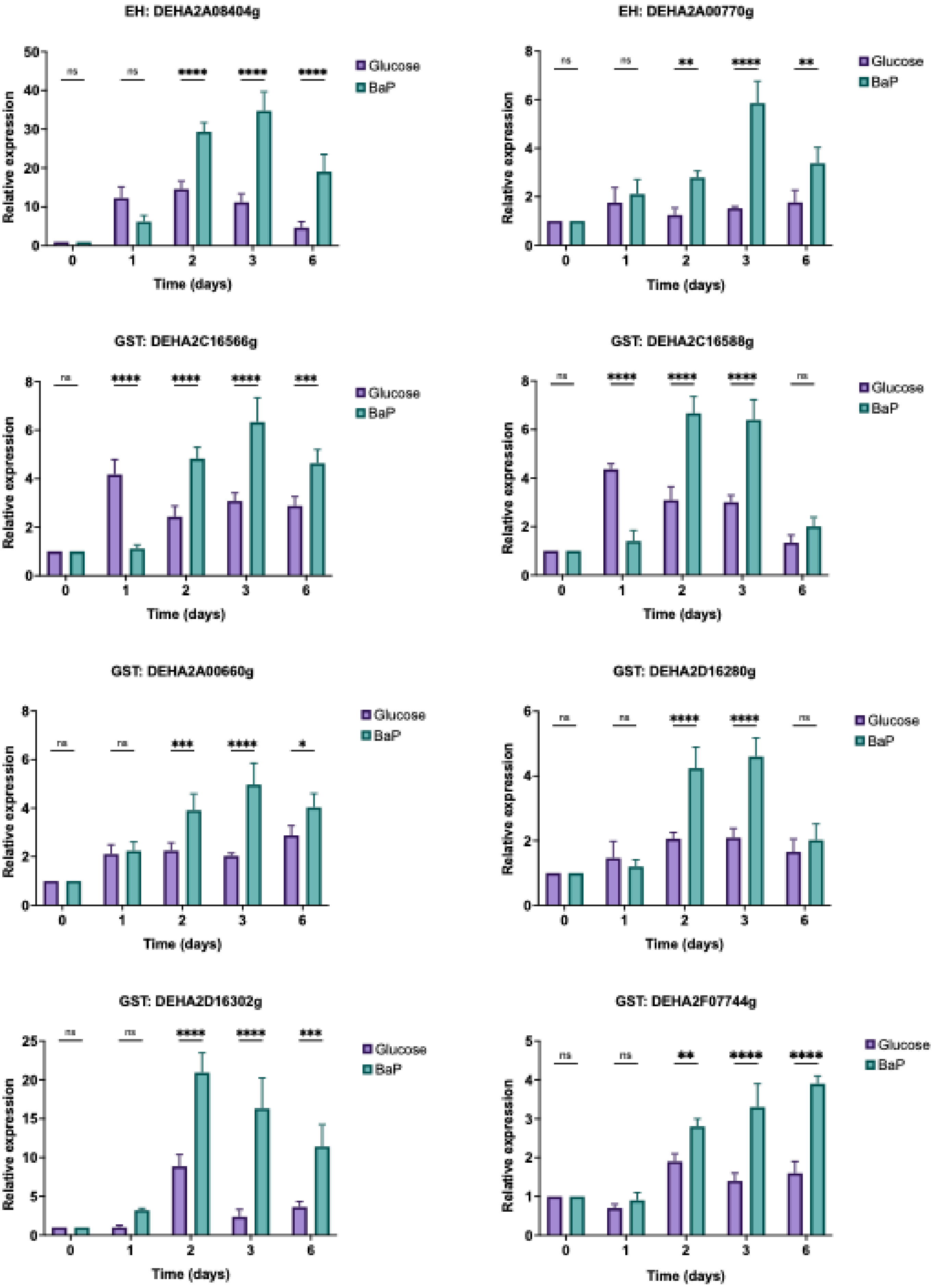
EH: DEHA2A08404g
Relative expression
Time (days)
Glucose
BaP
ns
ns
****
****
****
EH: DEHA2A00770g
Relative expression
Time (days)
Glucose
BaP
ns
ns
**
****
**
GST: DEHA2C16566g
Relative expression
Time (days)
Glucose
BaP
ns
****
****
****
***
GST: DEHA2C16588g
Relative expression
Time (days)
Glucose
BaP
ns
****
****
****
ns
GST: DEHA2A00660g
Relative expression
Time (days)
Glucose
BaP
ns
ns
***
****
*
GST: DEHA2D16280g
Relative expression
Time (days)
Glucose
BaP
ns
ns
****
****
ns
GST: DEHA2D16302g
Relative expression
Time (days)
Glucose
BaP
ns
ns
****
****
***
GST: DEHA2F07744g
Relative expression
Time (days)
Glucose
BaP
ns
ns
**
****
****

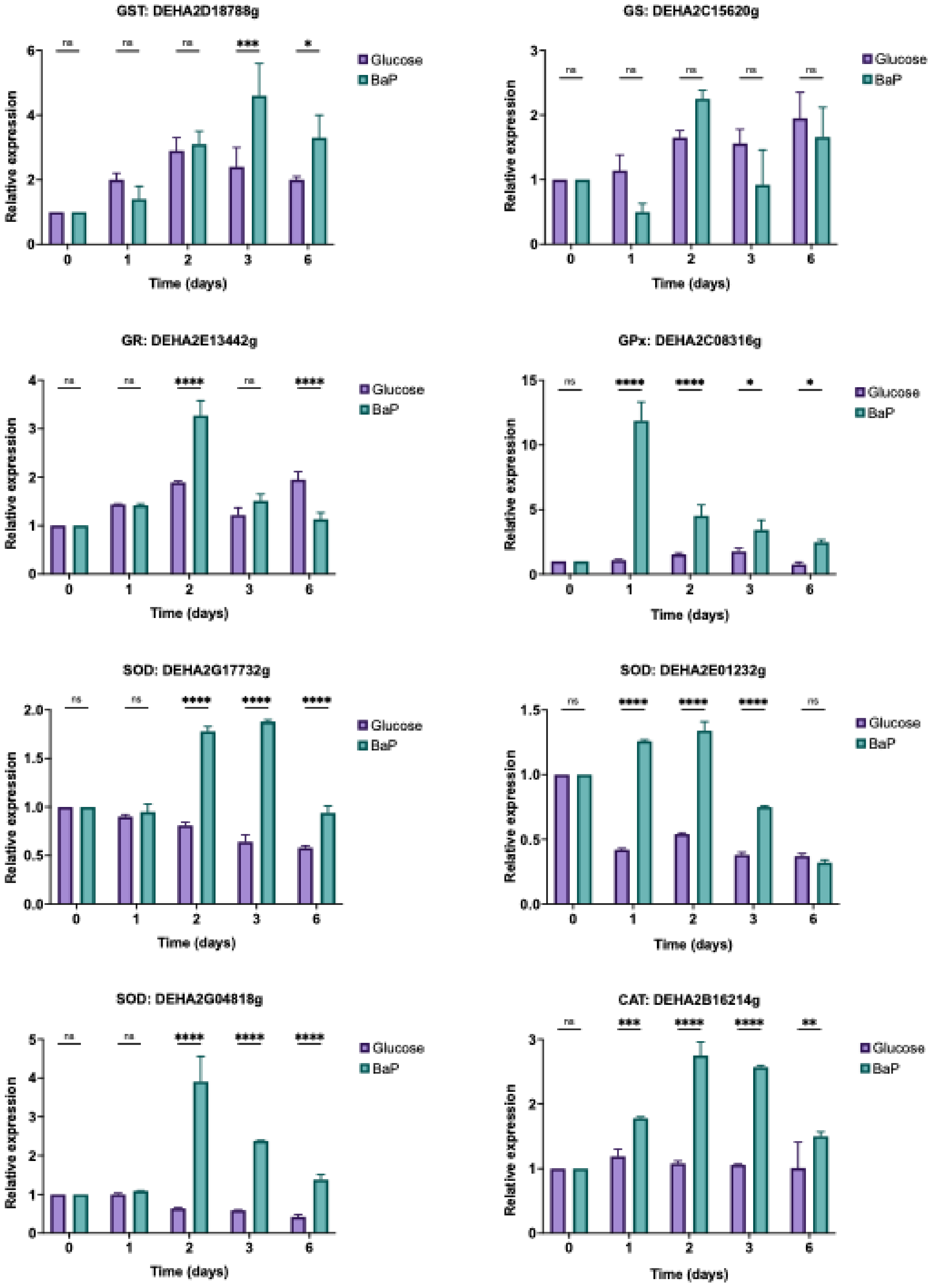


Data are shown as mean ± SD from four independent biological replicates, each measured in technical duplicate. Statistical significance was determined by two-way ANOVA with a full factorial model and Šidák's post hoc correction for multiple comparisons (α = 0.05). Adjusted $P$ values were reported as follows: ns ($P \geq 0.1234$), * ($P < 0.0332$), ** ($P < 0.0021$), *** ($P < 0.0002$), and **** ($P < 0.0001$).
